# A novel imaging ligand as a biomarker for mutant huntingtin-lowering in Huntington’s disease

**DOI:** 10.1101/2021.07.09.451725

**Authors:** Daniele Bertoglio, Jonathan Bard, Manuela Hessmann, Longbin Liu, Annette Gärtner, Stef De Lombaerde, Britta Huscher, Franziska Zajicek, Alan Miranda, Finn Peters, Frank Herrmann, Sabine Schaertl, Tamara Vasilkovska, Christopher J Brown, Peter D Johnson, Michael E Prime, Matthew R Mills, Annemie Van der Linden, Ladislav Mrzljak, Vinod Khetarpal, Yuchuan Wang, Deanna M Marchionini, Mette Skinbjerg, Jeroen Verhaeghe, Celia Dominguez, Steven Staelens, Ignacio Munoz-Sanjuan

## Abstract

Huntington’s disease (HD) is a dominantly inherited neurodegenerative disorder caused by a CAG trinucleotide expansion in the *huntingtin* (*HTT*) gene that encodes the pathologic mutant HTT (mHTT) protein with an expanded polyglutamine (PolyQ) tract. While several therapeutic programs targeting mHTT expression have advanced to clinical evaluation, no method is currently available to visualize mHTT levels in the living brain. Here we demonstrate the development of a positron emission tomography (PET) imaging radioligand with high affinity and selectivity for mHTT aggregates. This small molecule radiolabeled with ^11^C ([^11^C]CHDI-180R) enables non-invasive monitoring of mHTT pathology in the brain and can track region-and time-dependent suppression of mHTT in response to therapeutic interventions targeting mHTT expression. We further show that therapeutic agents that lower mHTT in the striatum have a functional restorative effect that can be measured by preservation of striatal imaging markers, enabling a translational path to assess the functional effect of mHTT lowering.

## Introduction

Neurodegenerative disease pathology is characterized by the presence of insoluble protein deposits in different subcellular compartments, which mark alterations in cellular homeostasis. Typically, neurodegenerative disorders have a complex molecular etiology, and affected brain cells display aggregation of a variety of proteins. In Huntington’s disease (HD), a CAG-tract expansion beyond 39 repeats in exon-1 of the *huntingtin* (*HTT*) gene is sufficient to cause the disease in a fully penetrant manner^1^. HD can be considered a multi-system atrophy disorder, even though the main pathological findings show ample degeneration of spiny projection neurons (SPNs) in the caudate and putamen, neurons in the globus pallidus and subthalamic nucleus of the basal ganglia, as well as significant but variable degeneration in neurons of the cerebral cortex and thalamic, cerebellar and hypothalamic nuclei^2^. Mutant huntingtin (mHTT) protein deposition in the neuropil and nucleus has variable morphology, is more frequent in some classes of projection neurons than in interneurons, and less frequent in cells of glial origin. A well-described progression “map” of degeneration pathology and aggregate deposition has been available for some time, although it is not clear how well histopathological changes inform the clinical staging of HD^2, 3^.

A longstanding goal for HD has been to target the cause of the disease. Therapeutic programs targeting HTT expression have advanced to clinical stages, including a now-terminated open-label extension study and Phase 3 trial that were evaluating the sustained safety and efficacy of tominersen, an antisense oligonucleotide (ASO) delivered intrathecally that can lower both mutant and wildtype (wt) HTT^4, 5^ (www.clinicaltrials.gov, identifier NCT03842969, NCT03761849, NCT03342053). The first gene therapy vector-mediated Phase 1/2 trial is now underway testing AMT-130, an AAV5-miRNA targeting both *HTT* alleles delivered directly into the caudate and putamen of HD patients^6^ (www.clinicaltrials.gov, identifier NCT04120493). Delivery of both of these agents is invasive and characterized by a restricted distribution that varies due to the modalities employed: the ASO predominantly decreases HTT expression in the spinal cord, cortical areas, and cerebellum, with some drugs reaching deeper basal ganglia nuclei, whereas the AAV-miRNA targets mostly the striatum and associated connected cell bodies via axonal transport^6, 7^. While the distribution and pharmacological activity of these therapeutics have been extensively evaluated in nonhuman primates and mHTT-expressing transgenic minipigs^7^ it is unclear whether we can expect a similar distribution in the larger human brain.

A key milestone was reached when the Ionis/Roche Phase 1/2a trial^4^ (www.clinicaltrials.gov, identifier NCT02519036) showed for the first time sustained dose- and time-dependent decreases in CSF levels of mHTT, demonstrating pharmacological activity in the human CNS. Regrettably, this finding has not led to clinical benefit in the recently-terminated Phase 3 tominersen trial, and analyses are underway to understand the safety issues identified, which led to a worsening of disease symptoms., How the reduction of mHTT in CSF after delivery of ASOs via lumbar puncture and AAVs delivered into brain parenchyma^5, 7^ relates to lowering in affected circuits in the brain is unclear.

To evaluate regional pharmacological effects of candidate therapeutics targeting mHTT, we sought to develop a non-invasive imaging agent specific for aggregated mHTT that could give insight into the timing, durability, and regional therapeutic effects of administered drugs^8, 9^. As all current therapeutic agents in development^10^ target either *HTT* or HTT transcriptional or post-transcriptional processes, quantification of mHTT protein offers a good indicator of the extent of HTT lowering and of the biodistribution of the agents.

For the first time, we here demonstrate the development of a PET imaging ligand with high affinity and selectivity for mHTT aggregation, that this polyQ-binding small molecule can detect mHTT aggregation in affected brain cells and can serve as a good indicator of pharmacological activity of agents that target HTT expression in the living brain. Specifically, we describe the ability of [^11^C]CHDI-180R, a nanomolar affinity small-molecule binder of aggregated, but not monomeric mHTT, to identify time-, dose- and region-specific pharmacological effects in two distinct interventional paradigms: direct striatal delivery of AAVs expressing ZFP repressors selectively targeting mHTT^11^ in the zQ175 HD mouse model^12, 13^, and in a novel genetically regulatable Q140 knock-in HD mouse model (the LacQ140^I^(*) model that enables ∼50% systemic lowering of m*Htt* mRNA and mHTT protein in a time-controlled manner. We show that [^11^C]CHDI-180R can accurately detect changes in mHTT levels early (within one month after m*Htt* lowering) and that the magnitude of suppression measured using [^11^C]CHDI-180R imaging correlates with mHTT levels quantified by immunoassays and classical histological evaluation. We further demonstrate that mHTT suppression can be measured after disease onset and that several imaging agents for striatal markers with diminished expression (PDE10 and dopamine receptors)^8, 14–18^ can detect the protective effects of mHTT lowering interventions in a time-dependent manner. We propose that imaging of PDE10a and dopamine receptors (D_1_R and D_2/3_R) can serve as functional response biomarkers for mHTT lowering with translational potential.

## Results

### CHDI-180 specifically binds mHTT in HD animal models

We show the ability of CHDI-180^8^, to detect aggregate pathology in HD mouse models, including the R6/2 (CAG 120) mice expressing an exon-1 mHTT protein^19^ and in the zQ175DN and HdhQ80 knock-in mouse models^12, 13, 20^.

Aggregate pathology was detected with [^3^H]CHDI-180 autoradiography (ARG) already at 4 weeks of age in R6/2 mice (Fig. 1a,b). In the zQ175DN heterozygous (het) model, aggregation is slower but detectable binding was measured at 6 months of age, increasing progressively until 13 months of age (Fig. 1c,d), in a pattern that mirrors histological analysis using mEM48 detection^21^. Since R6/2 and the zQ175DN models express large expansions in the polyQ tract, we explored the HdhQ80 KI model^20^ expressing smaller CAG lengths to understand if aggregate pathology could be detected in with finer temporal and spatial manner. Fig. 1e,f shows that CHDI-180 binding follows a ventro-dorsal gradient of aggregation within the striatum of HdhQ80 animals, beginning at 12m of age in the homozygotes (hom). (Fig. 1e,f). This pattern of aggregate pathology within the striatum was confirmed histochemically (Fig. 1g,h) with mEM48 antibody detection.

**Fig. 1.**
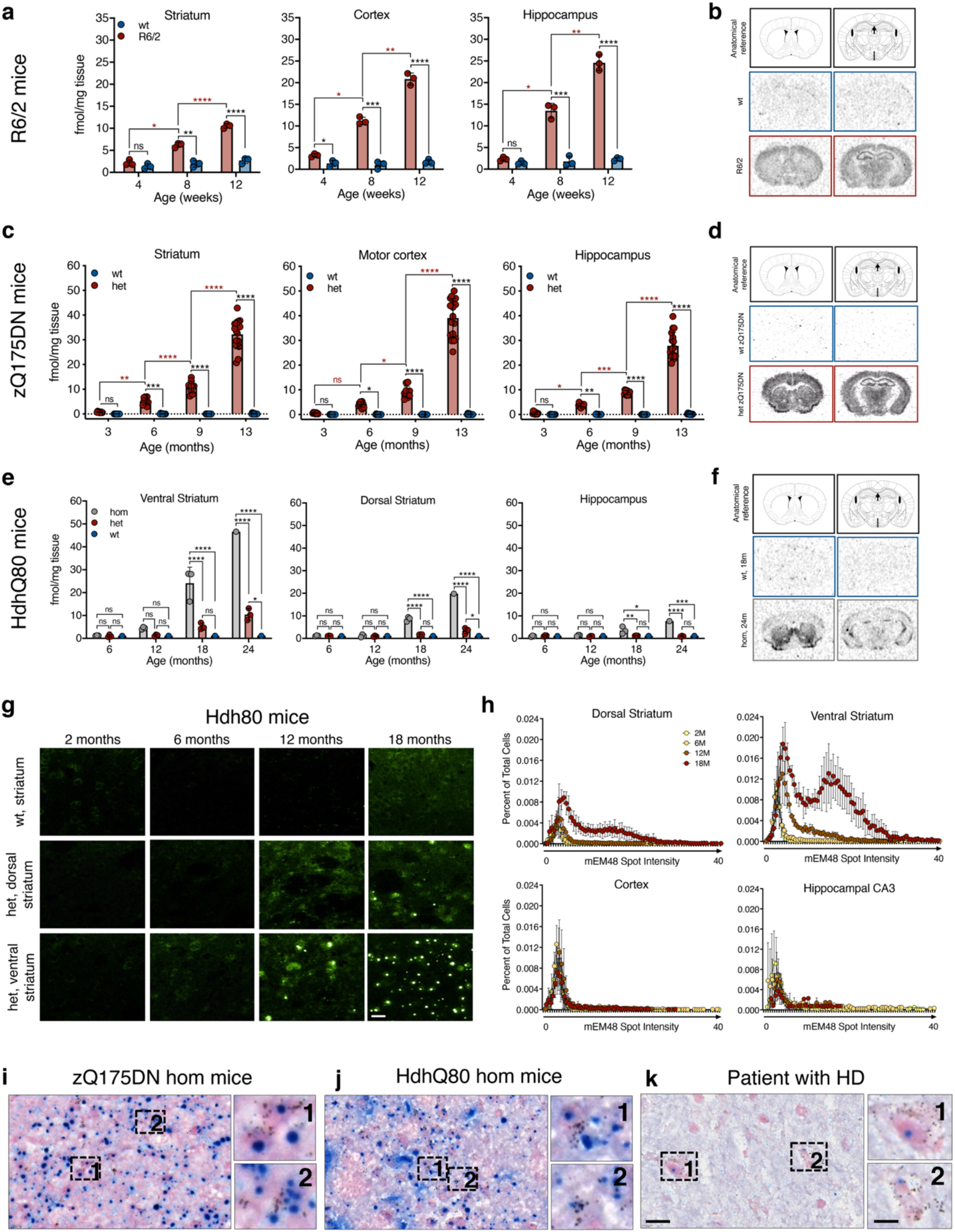
[^3^H]CHDI-180 mHTT-specific binding in HD mouse models without colocalizing with mHTT inclusions. **a-b**, Binding to transgenic R6/2 CAG120 mouse brains expressing mutant human exon1 Htt. **a**, Genotype-specific age-dependent increase in [³H]CHDI-180 in striatum, cortex, and hippocampus of 4-, 8-, and 12-week-old R6/2 CAG120 and wt littermates (wt, n = 3; R6/2, n = 3; per age). Two-way ANOVA with Tukey’s multiple comparison test; mean ± s.d., all points are shown; * *P* < 0.05, ** *P* < 0.01, *** *P* < 0.001, **** *P* < 0.0001. Red asterisk denotes signal differences between ages, as indicated for R6/2 mice. One representative study out of n > 15 experiments shown. **b**, Representative autoradiograms showing total binding of [³H]CHDI-180 in the striatum, cortex, and hippocampus of 12-week-old R6/2 and wt mice; anatomical orientation as indicated. **c-d**, Binding to knock-in zQ175DN het mouse brains carrying a humanized exon1 Htt sequence with 198 CAG repeats. **c**, Genotype-specific age-dependent increase in [³H]CHDI-180 in striatum, cortex, and hippocampus of 3, 6, 9 months (wt, n = 10; het, n = 10, per age), and 13 months (wt, n = 13; het, n = 17) of age. Two-way ANOVA with Tukey’s multiple comparison test; mean ± s.d., all points are shown; * *P* < 0.05, ** *P* < 0.01, *** *P* < 0.001, **** *P* < 0.0001. Red asterisk denotes signal differences between ages, as indicated for zQ175DN het mice. **d**, Representative autoradiograms showing total binding of [³H]CHDI-180 in striatum, cortex, and hippocampus of 13-month-old zQ175DN het and wt mice; anatomical orientation as indicated. **e-h**, Binding to knock-in HdhQ80 mouse brains carrying a humanized exon1 Htt sequence with 86 CAG repeats. **e**, Genotype-specific age-dependent increase in [³H]CHDI-180 in ventral and dorsal striatum as well as hippocampus at 6, 9, 18, and 24 months (wt, n = 1-3; het, n = 3; hom, n = 1-3, per age) of age. Two-way ANOVA with Tukey’s multiple comparison test; mean ± s.d., all points are shown; * *P* < 0.05, ** *P* < 0.01, *** *P* < 0.001, **** *P* < 0.0001. **f**, Representative autoradiograms showing total binding of [³H]CHDI-180 in striatum, cortex, and hippocampus of 24-month-old HdhQ80 hom and wt mice; anatomical orientation as indicated. **g**, Representative mHTT inclusions (mEM48) immunostaining in the dorsal and ventral striatum of HdhQ80 wt and het mice indicates that [³H]CHDI-180 binding is associated with the age- and brain region-dependent appearance of mEM48-positive mHTT inclusions as shown by mEM48 immunohistochemistry Scale bar, 20 *µ*m. **h**, Quantitative analysis of mEM48 intensity in HdhQ80 mice for mHTT inclusions in different brain regions and age groups. (**i-k**) Colocalization of [³H]CHDI-180 binding and mHTT inclusions (mEM48) in the ventral striatum of 12-month-old hom zQ175DN mice (**i**), ventral striatum of 24-month-old hom Hdh80 mice (**j**), and *post-mortem* frontal cortex of a patient with HD (#2017-060) (**k**). [³H]CHDI-180 silver grain signal was detectable in close vicinity to mEM48-positive signal but never co-registered with mHTT inclusion bodies, although it was partially co-registered with more diffuse appearing mEM48-positive signal. [^3^H]CHDI-180 binding, black silver grains; mHTT inclusions (mEM48), blue; background tissue (Nuclear Fast Red), pink. Scale bar, 20 *µ*m; inset, 10 *µ*m.

### CHDI-180 does not colocalize with nuclear inclusion bodies in HD animal models and human brains

The CHDI-180 ligand was initially identified using radioligand binding assays for expanded HTT proteins produced recombinantly^8^. However, mHTT aggregates come in different forms and can be detected in distinct subcellular compartments (intranuclear inclusions, diffuse nuclear-aggregated species, soma-localized aggregates, or neuropil aggregates). These species of oligomerized/aggregated mHTT can be detected with antibodies against aggregated, polyQ expanded mHTT, such as mEM48^22^ or PHP-1^23^. Therefore, we conducted double co-detection studies (binding and immunostaining), using [^3^H]CHDI-180 ARG and mEM48 immunohistochemistry (IHC) (Fig. 1i-j; Extended Data Fig. 1 and 2) or PHP-1 (not shown) in brain sections derived from zQ175DN and HdhQ80 mice and *post-mortem* human HD carriers. CHDI-180 binding did not co-localize with intranuclear inclusions detected by mEM48 in the zQ175DN model (Fig. 1i; Extended Data Fig. 2a-f) or in the HdhQ80 mouse model (Fig. 1j; Extended Data Fig. 1), with most signal observed outside the nucleus, presumably to neuropil or soma-localized mHTT aggregates. A similar pattern is observed in human brain samples from HD individuals (Fig. 1k; Extended Data Fig. 2g,h). There was no significant binding to the wt mouse brain nor in the brains of unaffected human subjects either in the grey or white matter under the autoradiographic conditions employed.

### PET imaging of mHTT pathology by [^11^C]CHDI-180R PET ligand

To examine the *in vivo* kinetic properties of [^11^C]CHDI-180R^8^ as a PET ligand, we selected the zQ175DN model because it displays a moderately slow disease onset with hallmark of mHTT-aggregates increasing from 3 to 12 months^21^. We performed *in vivo* microPET studies in 9-month-old zQ175DN het and wt mice for characterization of its pharmacokinetic properties and monitored its stability in the brain and plasma (Extended Data Fig. 3a).

Radio-high-performance liquid chromatography (radio-HPLC) coupled with γ-counter measurement of mouse brain homogenates and plasma samples did not show [^11^C]CHDI-180R-related metabolites in zQ175DN mice independent of genotype and mHTT inclusion levels (4- and 10-month-old het) (Extended Data Fig. 3b-c). Next, to evaluate [^11^C]CHDI-180R kinetics, we performed 90-min dynamic microPET scans following intravenous injection. We extracted an image-derived input function (IDIF) (Extended Data Fig. 3e) from the heart blood pool of each animal to serve as a non-invasive input function^24, 25^. Injection of [^11^C]CHDI-180R (Supplementary Table 1) resulted in a rapid radioactive uptake in the brain with standardized uptake value (SUV, regional radioactivity normalized to the injected activity and body weight) showing genotypic difference over the 90-min period and reversible kinetics described by a two-tissue compartment model (2TCM) (Extended data Fig. 3f, Supplementary Table 2). The resulting striatal total volume of distribution using IDIF (*V*_T (IDIF)_ as a surrogate of *V*_T_^26^) in het zQ175DN was significantly increased by 62% compared to wt littermates (Extended Data Fig. 3g, *P*<0.0001) with extremely low coefficients of variation (wt = 2.84%, het = 5.2%; Extended Data Fig. 3g). Scan acquisition could be reduced from 90-min down to 60-min (Extended Data Fig. 3h, R^2^ = 0.99, *P*<0.0001), and reliable *V*_T (IDIF)_ estimation of [^11^C]CHDI-180R binding was also obtained using the Logan graphical analysis^27^ as demonstrated by the optimal linear relationship (*y* = 1.08*x* - 0.04) with *V*_T (IDIF)_ estimation using 2TCM (Extended Data Fig. 3i, R^2^ = 0.99, *P*<0.0001). Finally, *V*_T (IDIF)_ parametric of [^11^C]CHDI-180R using the Logan model could be generated for both zQ175DN wt and het mice (Extended Data Fig. 3j).

### Longitudinal characterization of [^11^C]CHDI-180R PET ligand in zQ175DN mice

We performed a longitudinal evaluation of [^11^C]CHDI-180R microPET imaging in zQ175DN het and wt mice (Fig. 2a; Supplementary Table 3). In zQ175DN, mHTT-containing inclusions initiate in striatum^21^, and indeed the striatum was the first region where significant *V*_T (IDIF)_ differences were detected at 3 months of age (Fig. 2b,c). [^11^C]CHDI-180R *V*_T (IDIF)_ values revealed stable values over time in wt mice given the lack of specific target, while het zQ175DN displayed a significant temporal increase in all brain regions (e.g. in striatum, 6.7% (*P*<0.001), 40.3% (*P*<0.0001), 63.1% (*P*<0.0001), and 81.3% (*P*<0.0001), at 3, 6, 9 and 13 months of age, respectively) (Fig. 2c). For sample size requirements in therapeutic studies, see Supplementary Table 4. The increasing [^11^C]CHDI-180R binding within zQ175DN het was also confirmed by the voxel-based analysis of [^11^C]CHDI-180R *V*_T (IDIF)_ parametric maps, which could also identify specific cortical clusters of increased binding at advanced disease (13m > 9m) (Fig. 2d).

**Fig. 2.**
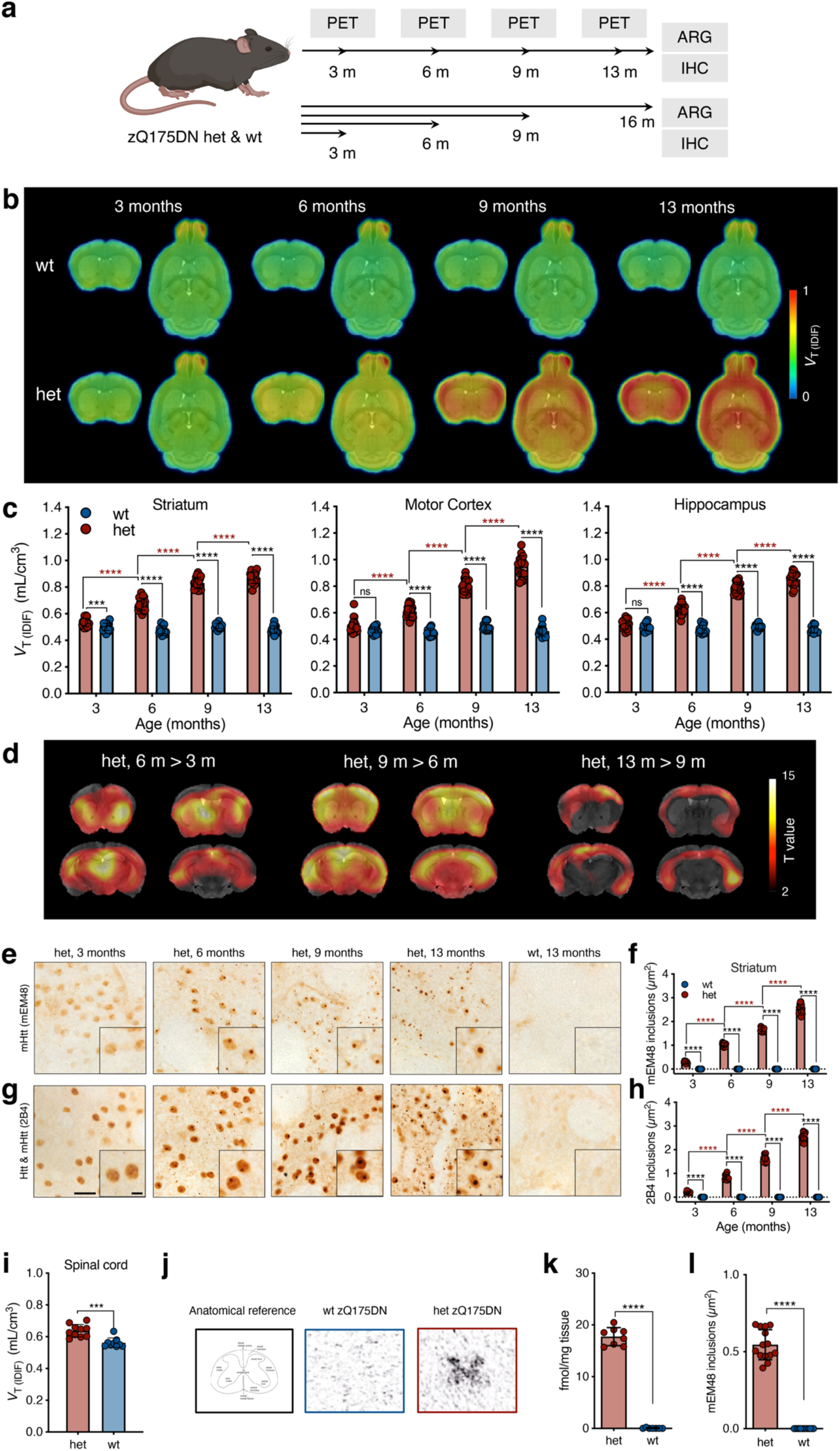
Longitudinal characterization of natural disease in zQ175DN het mice using [^11^C]CHDI-180R PET imaging. **a**, Timeline overview and endpoints in zQ175DN wt and het mice. **b**, Mean [^11^C]CHDI-180R *V*_T (IDIF)_ parametric images of zQ175DN wt and het mice at 3, 6, 9, and 13 months of age. PET images are co-registered to the MRI template for anatomical reference. Coronal and axial planes are shown. **c**, Regional [^11^C]CHDI-180R *V*_T (IDIF)_ quantification in zQ175DN wt and het at 3 months (wt, n = 19; het, n = 21), 6 months (wt, n = 15; het, n = 23), 9 months (wt, n = 13; het, n = 20), and 13 months (wt, n = 12; het, n = 17) of age. Repeated measures with linear mixed model analysis with Tukey-Kramer correction; *** *P* < 0.001, **** *P* < 0.0001. Data are shown as mean ± s.d., all points shown. Red asterisks denote longitudinal differences within zQ175DN het mice. **d**, Within zQ175DN het voxel-based analysis of [^11^C]CHDI-180R *V*_T (IDIF)_ parametric images. Comparison between 3-6 months (n = 21), 6-9 months (n = 20), and 9-13 months (n = 17) of age. Significant (*P* < 0.001) clusters are co-registered to the MRI template for anatomical reference and shown in the coronal panel. **e,g**, Genotype-specific age-dependent accumulation of mHTT inclusions in zQ175DN het mice at time points matching the longitudinal [^11^C]CHDI-180R PET study as demonstrated by mEM48 (**e**) and 2B4 (**g**) immunostaining. Scale bar, 20 *µ*m, inset scale bar, 5 *µ*m. **f,h**, Quantification of inclusions in wt and zQ175DN het mice for mEM48 (**f**) and 2B4 (**h**) at 3 months (wt, n = 10; het, n = 10), 6 months (wt, n = 10; het, n = 10), 9 months (wt, n = 10; het, n = 10), and 13 months (wt, n = 13; het, n = 17) of age. Two-way ANOVA with Bonferroni’s multiple comparison test; mean ± s.d., all points are shown; **** *P* < 0.0001. Red asterisks denote longitudinal differences within zQ175DN het mice. **i**, Spinal cord [^11^C]CHDI-180R *V*_T (IDIF)_ quantification in zQ175DN wt and het at 13 months (wt, n = 9; het, n = 10) of age. Two-tailed unpaired t-test with Welch’s correction; *** *P* < 0.001. Data are shown as mean ± s.d., all points shown. **j**, Representative autoradiograms showing total binding of [³H]CHDI-00485180 in the spinal cord of zQ175DN wt and het mice at 16 months; anatomical reference as indicated. **k**, Specific binding of [³H]CHDI-00485180 in the spinal cord of zQ175DN wt and het mice at 16 months (wt, n = 7; het, n = 8) of age. Two-tailed unpaired t-test with Welch’s correction; **** *P* < 0.0001. Data are shown as mean ± s.d., all points shown. **l**, Quantification of spinal cord inclusions in zQ175DN wt and het mice for mEM48 at 16 months (wt, n = 13; het, n = 15) of age. Two-tailed unpaired t-test with Welch’s correction; **** *P* < 0.0001. Data are shown as mean ± s.d., all points shown.

We monitored mHTT inclusions by mEM48 and 2B4^28^ immunoreactivity (Fig. 2a). In line with the [^11^C]CHDI-180R microPET findings, striatal mHTT inclusions could be observed starting at 3 months of age with a significant increase in size with disease progression in het zQ175DN mice for both mEM48 (Fig. 2e,f, *P*<0.0001) and 2B4^28^ (Fig. 2g,h, *P*<0.0001), while no mHTT inclusion was detected in wt littermates.

Within the CNS, mHTT inclusions are not limited to the brain as they may be found in the spinal cord in human patients, but this pathology has not been analyzed in mouse models of HD^29^. In the cervical spinal cord of zQ175DN het, [^11^C]CHDI-180R binding was significantly increased compared to wt littermates (Fig. 2i, *P*<0.001) as also confirmed by [^3^H]CHDI-180 ARG (Fig. 2j,k, *P*<0.0001) and mEM48 immunostaining (Fig. 2l, *P*<0.0001).

### [^11^C]CHDI-180R imaging identifies time- and region-dependent changes in mHTT pathology after virally-mediated, mHTT-selective striatal knockdown in the zQ175DN model

Given the ability of [^11^C]CHDI-180R to detect the temporal evolution of the mHTT pathology in live animals, we examined its applicability in measuring the effect of local or global mHTT lowering strategies (Fig. 3a, 4a, 5b; Supplementary Tables 5-9). We have previously demonstrated that striatal ZFP-mediated mHTT repression could improve molecular, histopathological, and electrophysiological deficits in the zQ175 het mice^11^. In this work, we used the ZFP-D repressor driven by the human synapsin promoter^11^ in two experimental paradigms to assess binding changes when the treatment is administered prior to disease onset (early treatment) versus after disease symptoms are well manifested (late treatment)^11–13^ (Fig. 3a, 4a). Het zQ175DN or wt mice were injected into striata with either AAV ZFP (treatment), ZFP-ΔDBD (ZFP lacking DNA-binding domain; control), or vehicle (PBS) before (2 months of age) or after (5 months of age) the age of mHTT inclusion formation and disease onset^12, 13^ (Fig. 3b, 4b). As shown in Figure 3b, we designed the experiment in a way that each animal acts as its own control; in one cohort, zQ175DN mice are injected with active ZFP in the left hemisphere, and with an inactive ZFP-ΔDBD in the right hemisphere. A second cohort of zQ175DN mice and a cohort of wild-type mice were injected with the vehicle in the left hemisphere, to control for the potential impact of viral transduction and exogenous protein expression, and with the control ZFP-ΔDBD in the right hemisphere. Animals were monitored longitudinally via [^11^C]CHDI-180R PET. In addition, other biomarkers known to undergo early, progressive, and profound changes years before clinical diagnosis, PDE10a, D_1_R, and D_2/3_R^14, 16, 30–32^, were assessed longitudinally using [^18^F]MNI-659, [^11^C]SCH23390, and [^11^C]Raclopride, respectively (Supplementary Tables 5,6). In the early intervention paradigm, mHTT pathology and PDE10a were assessed *in vivo*. During the late intervention paradigm, one study cohort was imaged for mHTT pathology and PDE10a, while a second cohort was analyzed for D_1_R and D_2/3_R. At the study end, *in vivo* findings were corroborated by ARG and immunostaining. Progressive alterations in these markers are recapitulated in zQ175DN het mice^8, 11, 15, 33–35^.

**Fig. 3.**
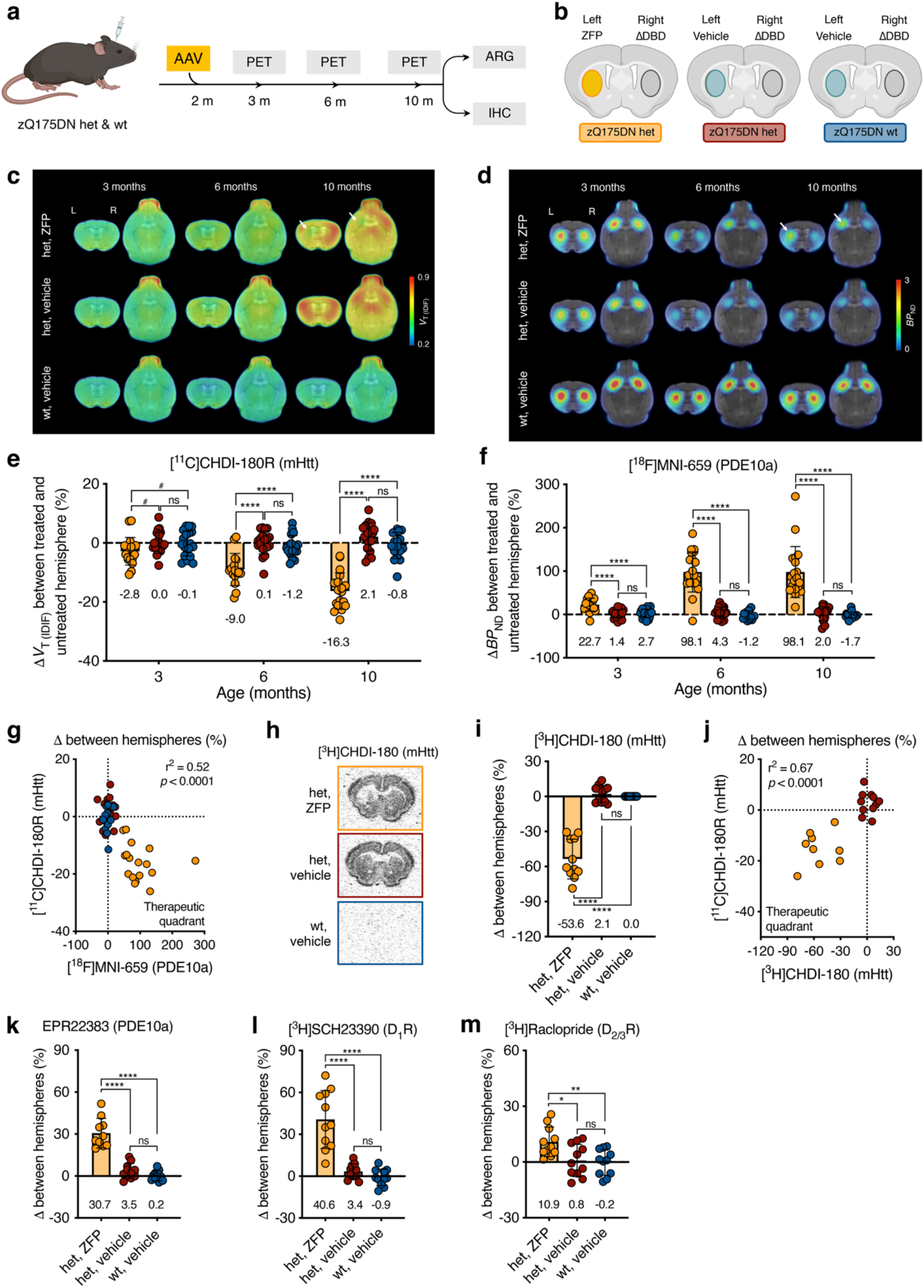
Response of [^11^C]CHDI-180R and imaging markers to early ZFP intervention in the striatum of zQ175DN mice. **a**, Timeline overview and endpoints of early ZFP intervention in zQ175DN wt and het mice. **b**, Experimental design overview in zQ175DN wt and het mice depicting injection hemisphere for ZFP treatment, ZFP-ΔDBD, and vehicle only. Fill colors represent the experimental group animals belong to. **c**,**d**, Mean [^11^C]CHDI-180R *V*_T (IDIF)_ (mHTT inclusions) (**c**) and [^18^F]MNI-659 *BP*_ND_ (PDE10a) (**d**) parametric images of zQ175DN wt vehicle, het vehicle, and het ZFP-treated mice at 3, 6, and 10 months of age. PET images are co-registered to the MRI template for anatomical reference. Coronal and axial planes are shown. A white arrow at 10 months of age indicates the ZFP-treated striatal hemisphere with reduced mHTT and increased PDE10a binding. **e**,**f**, Percentage contralateral difference for striatal [^11^C]CHDI-180R *V*_T (IDIF)_ (mHTT inclusions) (**e**) and [^18^F]MNI-659 *BP*_ND_ (PDE10a) (**f**) quantification in zQ175DN wt vehicle, het vehicle, and het ZFP-treated mice at 3, 6, and 10 months of age (het ZFP, n = 18-21; het vehicle, n = 18-22; wt vehicle, n = 18-20; values for each age, group, and radioligand) following striatal injection at 2 months of age. Repeated measures with linear mixed model analysis with Tukey-Kramer correction; ^#^ *P* < 0.1, **** *P* < 0.0001. Data are shown as mean ± s.d., all points shown. **g**, Correlation between contralateral difference for striatal mHTT and PDE10a binding with the het, ZFP group deviating from the center of axes towards the therapeutic quadrant. Two-tailed Pearson correlation analysis; R^2^ = 0.52; *P* < 0.0001. **h**, Representative autoradiograms showing total binding of [³H]CHDI-180 (mHTT inclusions) in zQ175DN wt vehicle, het vehicle, and het ZFP-treated mice. **i**, Percentage contralateral difference for striatal specific binding of [³H]CHDI-180 in zQ175DN wt vehicle, het vehicle, and het ZFP-treated mice at 10 months of age (het ZFP, n = 11; het vehicle, n = 11; wt vehicle, n = 11) following striatal injection at 2 months of age. One-way ANOVA with Tukey’s multiple comparison test; *** *P* < 0.001. Data are shown as mean ± s.d., all points shown. **j**, Correlation between contralateral difference for striatal mHTT binding measured with microPET and autoradiography at 10 months of age depicting the het ZFP-treated mice deviating from the center of axes towards the therapeutic quadrant. Two-tailed Pearson correlation analysis; R^2^ = 0.67; *P* < 0.0001. **k-m**, Percentage contralateral difference for PDE10a immunostaining (**k**), [³H]SCH23390 (D_1_R) (**l**), [³H]Raclopride (D_2/3_R) (**m**) in zQ175DN wt vehicle, het vehicle, and het ZFP-treated mice at 10 months of age (het ZFP, n = 11; het vehicle, n = 11; wt vehicle, n = 11) following striatal injection at 2 months of age. One-way ANOVA with Tukey’s multiple comparison test; * *P* < 0.05, ** *P* < 0.01, **** *P* < 0.0001. Data are shown as mean ± s.d., all points shown.

**Fig. 4.**
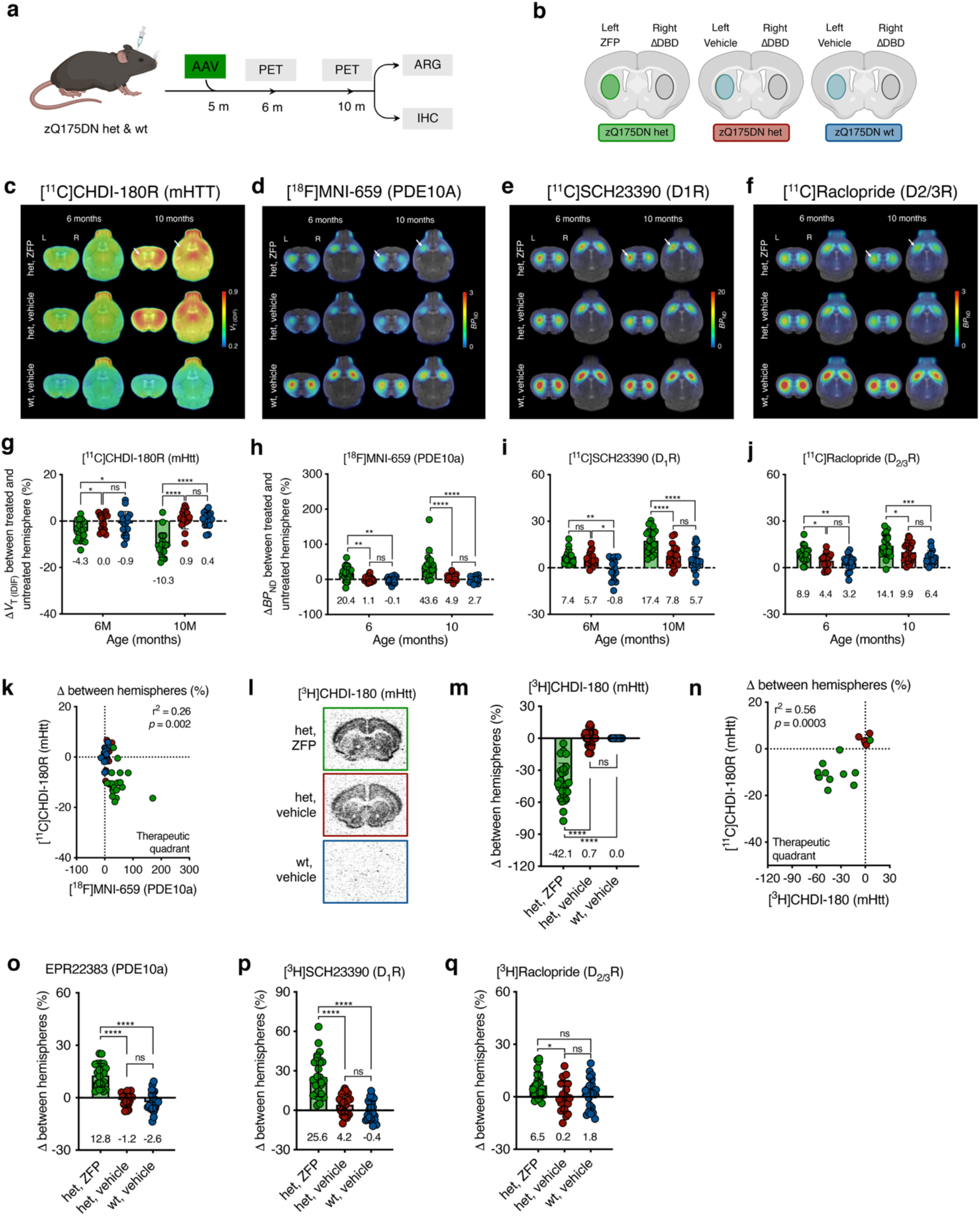
Response of [^11^C]CHDI-180R and imaging markers to late ZFP intervention in the striatum of zQ175DN mice. **a**, Timeline overview and endpoints of late ZFP intervention in zQ175DN wt and het mice. **b**, Experimental design overview in zQ175DN wt and het mice depicting injection hemisphere for ZFP treatment, ZFP-ΔDBD, and vehicle only. Fill colors represent the experimental group animals belong to. **c**-**f**, Mean [^11^C]CHDI-180R *V*_T (IDIF)_ (mHTT inclusions) (**c**), [^18^F]MNI-659 *BP*_ND_ (PDE10a) (**d**), [^11^C]SCH23390 *BP*_ND_ (D_1_R) (**e**), and [^11^C]Raclopride *BP*_ND_ (D_2/3_R) (**f**) parametric images of zQ175DN wt vehicle, het vehicle, and het ZFP-treated mice at 6 and 10 months of age. PET images are co-registered to the MRI template for anatomical reference. Coronal and axial planes are shown. A white arrow at 10 months of age indicates the ZFP-treated striatal hemisphere with reduced mHTT as well as increased PDE10a, D_1_R, and D_2/3_R binding. **g-j**, Percentage contralateral difference for striatal [^11^C]CHDI-180R *V*_T (IDIF)_ (mHTT inclusions) (**g**), [^18^F]MNI-659 *BP*_ND_ (PDE10a) (**h**), [^11^C]SCH23390 *BP*_ND_ (D_1_R) (**i**), and [^11^C]Raclopride *BP*_ND_ (D_2/3_R) (**j**) quantification in zQ175DN wt vehicle, het vehicle, and het ZFP-treated mice at 6 and 10 months of age (het ZFP, n = 17-23; het vehicle, n = 16-22; wt vehicle, n = 16-19; values for each age, group, and radioligand) following striatal injection at 5 months of age. Repeated measures with linear mixed model analysis with Tukey-Kramer correction; * *P* < 0.05, ** *P* < 0.01, *** *P* < 0.001, **** *P* < 0.0001. Data are shown as mean ± s.d., all points shown. **k**, Correlation between contralateral difference for striatal mHTT and PDE10a binding with the het, ZFP group partly deviating from the center of axes towards the therapeutic quadrant. Two-tailed Pearson correlation analysis; R^2^ = 0.26; *P* = 0.002. **l**, Representative autoradiograms showing total binding of [³H]CHDI-180 (mHTT inclusions) in zQ175DN wt vehicle, het vehicle, and het ZFP-treated. **m**, Percentage contralateral difference for striatal specific binding of [³H]CHDI-180 in zQ175DN wt vehicle, het vehicle, and het ZFP-treated mice at 10 months of age (het ZFP, n = 25; het vehicle, n = 26; wt vehicle, n = 25) following striatal injection at 5 months of age. One-way ANOVA with Tukey’s multiple comparison test; **** *P* < 0.0001. Data are shown as mean ± s.d., all points shown. **n**, Correlation between contralateral difference for striatal mHTT binding measured with microPET and autoradiography at 10 months of age depicting the het ZFP-treated mice partly deviating from the center of axes towards the therapeutic quadrant. Two-tailed Pearson correlation analysis; R^2^ = 0.56; *P* = 0.0003. **o-q**, Percentage contralateral difference for immunostaining for PDE10a (**o**), and autoradiography for [³H]SCH23390 (D_1_R) (**p**), [³H]Raclopride (D_2/3_R) (**q**) in zQ175DN wt vehicle, het vehicle, and het ZFP-treated mice at 10 months of age (het ZFP, n = 25; het vehicle, n = 26; wt vehicle, n = 25) following striatal injection at 5 months of age. One-way ANOVA with Tukey’s multiple comparison test; * *P* < 0.05, **** *P* < 0.0001. Data are shown as mean ± s.d., all points shown.

**Fig. 5.**
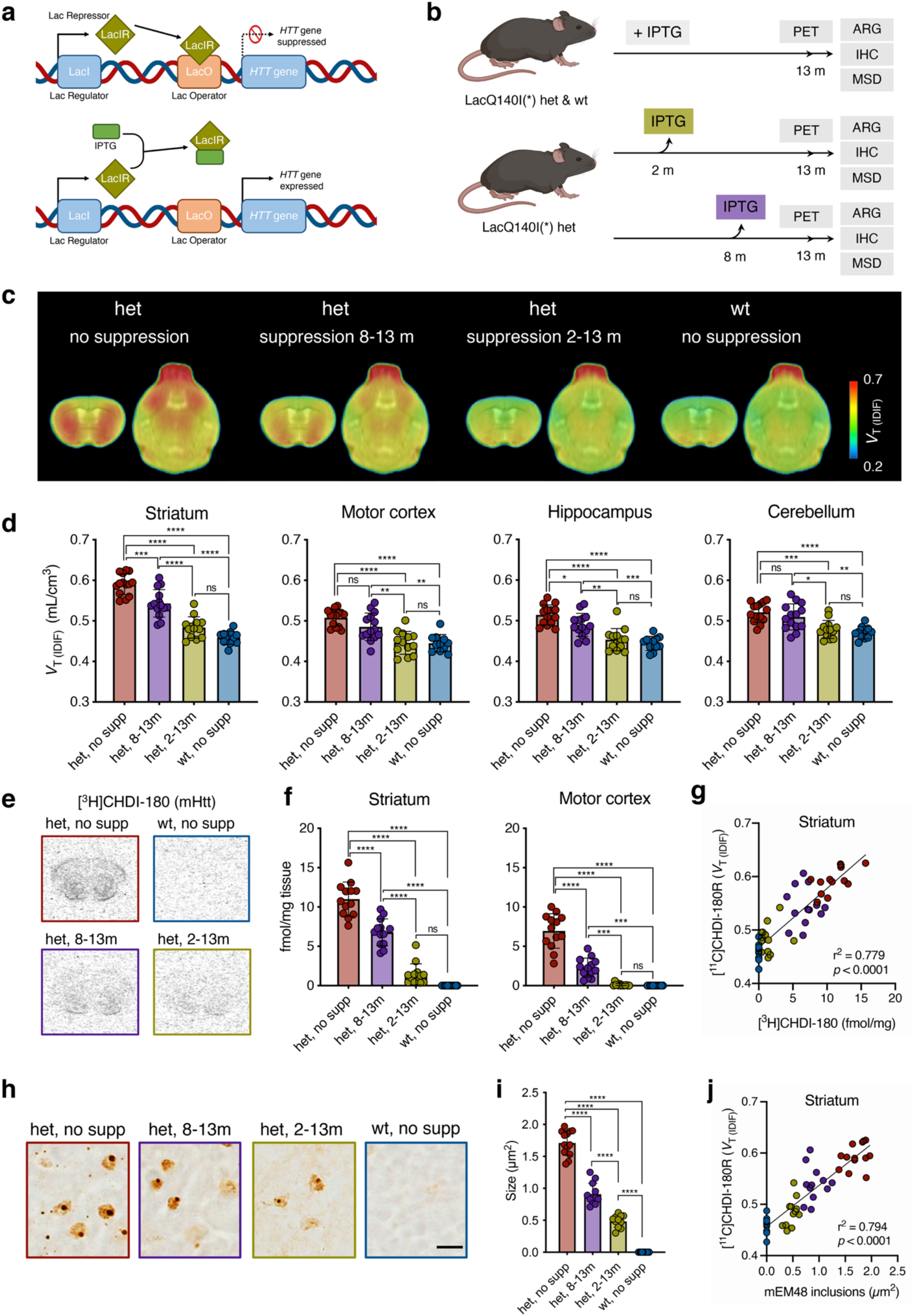
Modulation of [^11^C]CHDI-180R binding by broadly distributed mHTT lowering in LacQ140^I^(*) het mice. **a**, Schematic overview of the LacQ140^I^(*) allele. The transcriptional repressor, LacIR, binds to the Lac Operator, LacO, precluding expression of the Q140 allele (top). Administration of isopropyl β-d-1-thiogalactopyranoside, IPTG, allosterically inhibits LacIR, allowing transcription of the Q140 allele. **b**, Timeline overview and endpoints in LacQ140^I^(*) wt and het mice. **c**, Mean [^11^C]CHDI-180R *V*_T (IDIF)_ parametric images of LacQ140^I^(*) wt and het mice at 13 months of age. PET images are co-registered to the MRI template for anatomical reference. Coronal and axial planes are shown. **d**, Regional [^11^C]CHDI-180R *V*_T (IDIF)_ quantification in LacQ140^I^(*) wt and het at 13 months (het no supp, n = 13; het 8-13m, n = 14; het 2-13m, n = 12; wt no supp, n = 13) of age. One-way ANOVA with Tukey’s multiple comparison test; * *P* < 0.05, ** *P* < 0.01, *** *P* < 0.001, **** *P* < 0.0001. Data are shown as mean ± s.d., all points shown. **e**,**h**, Representative autoradiograms showing total binding of [³H]CHDI-00485180 (mHTT inclusions) (**e**) and immunostaining as demonstrated by mEM48 (**h**) in LacQ140^I^(*) wt and het mice. Scale bar, 10 *µ*m. **f**,**i**, Specific binding of [³H]CHDI-00485180 (**f**) and quantification of inclusions for mEM48 (**i**) in LacQ140^I^(*) wt and het mice at 13 months of age (het no supp, n = 13; het 8-13m, n = 14; het 2-13m, n = 12; wt no supp, n = 13). One-way ANOVA with Tukey’s multiple comparison test; *** *P* < 0.001, **** *P* < 0.0001. Data are shown as mean ± s.d., all points shown. **g**, Correlation between striatal mHTT binding measured with microPET and autoradiography in LacQ140^I^(*) wt and het mice at 13 months of age. Two-tailed Pearson correlation analysis; R^2^ = 0.779; *P* < 0.0001. **j**, Correlation between striatal mHTT binding measured with microPET and immunostaining in LacQ140^I^(*) wt and het mice at 13 months of age. Two-tailed Pearson correlation analysis; R^2^ = 0.794; *P* < 0.0001.

In the early ZFP intervention (Fig. 3a,b; Supplementary Tables 5 and 8), [^11^C]CHDI-180R *V*_T (IDIF)_ values for the ZFP-versus ΔDBD-injected striatum were significantly reduced by 2.8, 9.0, and 16.3% at 3, 6, and 10 months of age, respectively (Fig. 3c, e; treatment effect: *P*<0.0001). No difference was observed for control zQ175DN cohorts (Fig. 3c,e; Extended Data Fig. 4a,b). The reduced [^11^C]CHDI-180R binding was paralleled by a significant increase in the non-displaceable binding potential (*BP*_ND_, a quantitative index of receptor density^26^) for [^18^F]MNI-659 (22.7%, 98.1%, and 98.1% at 3, 6, and 10 months of age, respectively, treatment effect: *P*<0.0001; or 1-, 4- and 8 months after viral transduction with ZFP-mediated mHTT suppression), with no contralateral difference for the control zQ175DN cohorts (Fig. 3d,f; Extended Data Fig. 4c,d). The estimated “therapeutic” effect for the early intervention, calculated according to eq. 1, suggested approximately 40% mHTT lowering (42.2, 40.6, and 38.8% at 3, 6, and 10 months of age), which was positively associated with the 43.3% PDE10a preservation in the same animals (Fig. 3g; R^2^ = 0.52, *P*<0.0001). Upon completion of the studies, ARG was performed using [^3^H]CHDI-180 (mHTT), [^3^H]SCH23390 (D_1_R), and [^3^H]Raclopride (D_2/3_R) as well as immunostaining for PDE10a. Striatal [^3^H]CHDI-180 binding for the ZFP-versus ΔDBD-injected striatum was significantly reduced by 53.6% (Fig. 3h,i, *P*<0.0001), showing correlation with the *in vivo* [^11^C]CHDI-180R PET measurement (Fig. 3j; R^2^ = 0.67, *P*<0.0001). A significant increase in ZFP-versus ΔDBD-injected striatum was measured for PDE10a immunostaining (30.7%, *F*_(2,30)_ = 59.40, *P*<0.0001), D_1_R with [^3^H]SCH23390 (40.6%, *F*_(2,30)_ = 34.98, *P*<0.0001), and D_2/3_R with [^3^H]Raclopride (10.9%, *F*_(2,30)_ = 6.59, *P*<0.01) (Fig. 3k-m). Noteworthy, the reduction in mHTT levels was correlated with preservation of all measured striatal markers (PDE10a: R^2^ = 0.84, *P*<0.0001; D_1_R: R^2^ = 0.79, *P*<0.0001; D_2/3_R: R^2^ = 0.29, *P*=0.0012; Extended Data Fig. 5).

In the late ZFP intervention paradigm (Fig. 4a,b; Supplementary Tables 6 and 9), [^11^C]CHDI-180R *V*_T (IDIF)_ values for the ZFP-versus ΔDBD-injected striatum were significantly reduced by 4.3% and 10.3% at 6 and 10 months of age (*P*<0.0001, 1 month and 5 months post viral transduction), respectively, without contralateral differences for control zQ175DN cohorts (Fig. 4c,g; Extended Data Fig. 6a,b). In addition, a significant increase in *BP*_ND_ for ZFP-compared to ΔDBD-ZFP injected striatum was measured for all translational biomarkers, with [^18^F]MNI-659 being increased by 20.4% and 43.6% (Fig. 4d, h; Extended Data Fig. 6c,d), [^11^C]SCH23390 by 7.4% and 17.4% (Fig. 4e, i; Extended Data Fig. 6e,f), and [^11^C]Raclopride by 8.9% and 14.1% (Fig. 4f, j; Extended Data Fig. 6g,h) at 6 and 10 months, respectively (treatment effect: *P*<0.0001 for all markers). However, when the percentage difference in *BP*_ND_ between hemisphere was corrected by the het control group, the ZFP group displayed increased binding of 19.3% and 38.7% ([^18^F]MNI-659), by 1.7% and 9.6% ([^11^C]SCH23390), and by 4.5% and 4.2% ([^11^C]Raclopride) at 6 and 10 months, respectively (Fig. 4h-j, treatment effect: *P*<0.0001 for all markers). The estimated therapeutic effect for the late intervention, (eq. 1), indicated approximately 23% mHTT lowering (19.4% and 23.6% at 6 and 10 months of age), positively associated to the 25.5% PDE10a preservation (Fig. 4k; R^2^ = 0.26, *P*=0.002). In *post-mortem* experiments, [^3^H]CHDI-180 binding for the ZFP-versus ΔDBD-injected striatum was significantly reduced by 42.1% (Fig. 4l,m), showing agreement with [^11^C]CHDI-180R (Fig. 4n; R^2^ = 0.56, *P*=0.0003). Notably, this effect was lower than the 53.6% measured during early intervention (Fig. 3i), possibly due to time of intervention (2 or 5 months of age), the duration of the treatment (8 months or 5 months), or a combination of these factors. Additionally, we observed a significant increase in ZFP-versus ΔDBD-injected striatum for PDE10a (12.8%, *F*_(2,63)_ = 52.75, *P*<0.0001), D_1_R with [^3^H]SCH23390 (25.6%, *F*_(2,66)_ = 37.70, *P*<0.0001), and D_2/3_R with [^3^H]Raclopride (6.5%, *F*_(2,66)_ = 3.71, *P*=0.0297) (Fig. 4o-q). The preservation of all measured striatal markers in the ZFP-injected hemisphere was correlated with the reduction in mHTT levels (PDE10a: R^2^ = 0.61, *P*<0.0001; D_1_R: R^2^ = 0.48, *P*<0.0001; D_2/3_R: R^2^ = 0.14, *P*=0.0016; Extended Data Fig. 5).

Meso Scale Discovery (MSD) measurements showed that ZFP treatment did not alter wt mouse HTT levels, but significantly decreased levels of soluble and aggregated mHTT in both the early (39% and 69%, respectively; *P*<0.0001, Extended Data Fig. 7a-c) and late (43% and 37%, respectively; *P*<0.05, Extended Data Fig. 7d-f) intervention studies. Soluble expanded HTT protein was detected using 2B7-MW1 assay^36^ and aggregated mHTT was detected using MW8-4C9 assay^37^. Most likely, mHTT reduction on a level of only ZFP expressing cells would be higher since we measured an average of 38% AAV-ZFP transduced cells along the rostro-caudal axis, following AAV ZFP and ΔDBD-ZFP injections (Extended Data Fig. 8). We analyzed several brains to confirm the extent of ZFP distribution and its impact on mHTT aggregate number and intensity using mEM48 immunohistochemistry. In the early treatment paradigm, the striatal region expressing the AAV ZFP treatment did not display any mHTT nuclear inclusions at 10 months of age (8 months of treatment) unlike the ZFP untransduced region or contralateral ΔDBD-ZFP injected hemisphere (Extended Data Fig. 9a,b). In contrast, in the late treatment paradigm, smaller and fewer intranuclear mHTT inclusions are present following the AAV ZFP treatment than in the ZFP untransduced area or the contralateral ΔDBD-ZFP injected hemisphere (Extended Data Fig. 9c,d). No evidence of microglial or astrocytic reactivity in the striatum of injected animals, as judged by Iba1 and GFAP reactivity (Extended Fig. 10), recapitulating what we reported in Zeitler et al^11^.

Collectively, these observations suggest that we achieve a 40% mHTT reduction in the early intervention paradigm and that this value is determined by the extent of neuronal transduction and viral distribution in the mouse striatum (38%). This value is in concordance with the 38.9% signal decrease, measured *in vivo* using PET, with [^11^C]CHDI-180R. In contrast, in the late intervention paradigm, and consistent with residual pre-existing aggregate pathology that remains after AAV-ZFP administration after disease onset, we only achieved a 23.6% therapeutic effect as measured *in vivo* with [^11^C]CHDI-180R at 10 months of age.

### [^11^C]CHDI-180R imaging detects widespread suppression of mHTT in a novel, regulatable mHTT knock-in LacQ140^I^(*) mouse model

Current clinical HTT-lowering directed therapeutic strategies are attempting to lower HTT by 50% in cortical and striatal regions, depending on the modality^5, 6, 10, 38, 39^. Therefore, we wanted to detect CNS-wide changes in mHTT within the range being pursued clinically, using a newly characterized knock-in LacQ140^I^(*) mouse model, which allows for m*Htt* lowering in a regulatable fashion to approximately 40-50% throughout the body in a Q140 KI context^40^. Due to the presence of the LacO repressor binding sites, the exposure to IPTG (isopropyl-β-d-1-thiogalactopyranoside) enables (i.e. derepresses) the expression of mHTT. Upon withdrawal of IPTG, mHTT expression is suppressed throughout the organism (Fig. 5; Extended Data Fig. 7g-i). The extent of mHTT aggregated species, as judged by MSD assays with MW8-4C9 (Extended fig. 7h) and histological analysis (not shown) depends on the timing of mHTT mRNA suppression.

We employed this model to lower m*Htt* systemically at 2 or 8 months of age, before and after mHTT inclusion formation and disease onset, and compared them to control mice or LacQ140^I^(*) mice with m*Htt* expressed throughout its life, at 13 months of age (Fig. 5a,b; Supplementary Tables 7). [^11^C]CHDI-180R *V*_T (IDIF)_ values were significantly reduced consistently with suppression duration in all brain regions examined following IPTG withdrawal before (2-13 months) and after (8-13 months) mHTT inclusion formation (Fig. 5c,d). The estimated mHTT suppression effect, calculated according to eq. 2 in methods, suggested global 80-95% or 20-35% mHTT aggregate lowering following IPTG withdrawal at 2 or 8 months (treatment effect striatum: *F*_(3,48)_ = 62.30, *P*<0.0001).

Consistently, autoradiographic [^3^H]CHDI-180 binding was significantly reduced (Fig. 5e,f; striatum, *F*_(3,47)_ = 133.80, *P*<0.0001) demonstrating agreement with [^11^C]CHDI-180R PET (Fig. 5g, R^2^ = 0.779, *P*<0.0001). The extent of mHTT lowering was supported by mEM48 immunostaining (Fig. 5h,i; *F*_(3,46)_ = 360.80, *P*<0.0001), in line with the [^11^C]CHDI-180R binding (Fig. 5j, R^2^ = 0.794, *P*<0.0001), as well as MSD measurements of HTT using cerebellar extracts obtained from the same animals (Extended Data Fig. 7g-i; *P*<0.05-*P*<0.0001).

## Discussion

Therapeutic studies targeting HTT expression with ASOs and AAV-miRNAs being evaluated or planned in clinical studies^5, 6, 10, 38, 39^. Given the different therapeutic modalities leading to distinct restricted distribution patterns, an understanding of the regional effects of HTT lowering agents is fundamental in being able to interpret, and improve upon, clinical trial results. It is in this context that we set out to develop a translational biomarker strategy to identify and characterize potential biomarkers that can help guide the clinical development of HTT lowering agents. Here we extend our prior characterization of CHDI-180 and demonstrate the time- and region-dependent appearance of mHTT pathology in the HD mouse models R6/2, HdhQ80, and zQ175DN. The ligand is suitable to detect genotype and region-specific differences in HTT pathology throughout the brain, allowing for its deployment in therapeutic studies with manageable sample size and a longitudinal manner. We were able to ascertain different regional pathology within the striatum, particularly in HdhQ80 mice, which appears to proceed from a ventral to dorsal trajectory, an observation reminiscent of human pathology that proceeds caudal-to-rostral and dorsal-to-ventral^2, 3^.

We applied [^11^C]CHDI-180R in two interventional paradigms when mHTT is lowered in a restricted manner in the striatum of mice, or more broadly throughout the mouse brain, within the range of mHTT suppression expected in clinical studies (∼50%). The extent of lowering detected by [^11^C]CHDI-180R correlates well with the extent of mHTT suppression as measured by quantitative assays for soluble and aggregated forms of mHTT. These studies show that [^11^C]CHDI-180R can be used irrespective of the regional distribution of the therapeutic agents or the extent of lowering. Furthermore, we verified the extent of lowering by ARG, showing excellent concordance with PET imaging. In the context of the ZFP repressor, the decrease in signals obtained with [^11^C]CHDI-180R appear rapid (1-month post administration of AAV-ZFP), and are sustained during the duration of the studies (up to 8 months). When administered early, prior to the appearance of pathology, AAV-ZFP prevents mHTT inclusion and extranuclear aggregation, and the diminution of the signal detected by [^11^C]CHDI-180R can be explained by the extent of agent distribution and neuronal transduction (in our case, about 40% of the striatum).

While we do not yet have a full understanding of the various species of mHTT that constitute the binding site(s) for CHDI-180, based on recombinant, cell, and tissue protein studies^8, 9^, we know this ligand can bind oligomerized and some forms of fibrillar mHTT but not to monomeric soluble HTT.

We also investigated potential striatal markers that can serve as markers of functional SPN restoration. Several PET ligands, previously shown to track disease progression in HD individuals, have been shown to track progression in models of HD^33–35, 41, 42^. We show that the response to mHTT lowering in SPNs is fast and durable, and that these effects can be observed even in the context of established disease and aging, at least in the zQ175DN model. During the early intervention paradigm, all striatal markers responded within a month of therapy administration, suggesting an improvement of cellular alterations in indirect-pathway SPNs (expressing both PDE10 and D_2_R), and direct-pathway neurons (expressing PDE10 and D_1_R). When AAV-ZFP is administered after disease onset, the response is more muted, but significant for all tracers, particularly as judged by ARG, which has a higher signal to background ratio than microPET. PDE10a and D_1_R expression appear more responsive to mHTT lowering than D_2/3_R, arguing that direct pathway neurons (affected later in the disease) might be more amenable to functional restoration.

The strong correlation seen in intra-animal comparisons between [^11^C]CHDI-180R and PDE10a binding across our cohorts strongly support the concept that PDE10a imaging can be a very sensitive translational marker of mHTT lowering. As this marker is one of the earliest markers altered in premanifest individuals, including those far from disease onset^14–16^, PDE10a imaging can be used to track functional responses to HTT lowering in prodromal clinical studies.

In summary, we demonstrated the development of a small-molecule PET ligand with high affinity and selectivity for mHTT to monitor non-invasively mHTT pathology in the living brain and track region- and time-dependent suppression of mHTT levels in response to therapeutic intervention. We also showed that therapeutic agents, such as AAV-ZFP, can be functionally restorative and their effects can be measured by the preservation of striatal imaging markers.

## Materials and Methods

### *In vitro* studies using IHC and ARG

#### Brain preparation mouse models of HD and human brain tissue

For autoradiography (ARG) and immunohistochemistry (IHC) analysis, fresh frozen whole brain samples from male zQ175DN, HdhQ80, and R6/2 and age-matched wt mice were prepared (Supplementary Table 10). For ARG analyses, mice were either euthanized by cervical dislocation or PBS-perfusion and the brains were snap-frozen in isopentane at −30 to −40°C and stored at −80°C. For IHC analyses, the mice were euthanized by PBS-perfusion followed by 4% paraformaldehyde (PFA). The brains were fixed in 4% paraformaldehyde (PFA) fixation for 24 hours, followed by 48 hours in 30% sucrose at 4°C. Afterwards the brains were embedded in Tissue-Tek® O.C.T. Compound (Sakura, Cat. # 4583) using Peel-A-Way^TM^ embedding molds (Sigma Aldrich, Cat. # E6032-1CS) and stored at −80°C.

For ARG and IHC analysis in human brain tissue, *post-mortem* superior frontal gyrus tissue from HD patients and control donors without any evidence of neurological disease were obtained from Netherlands Brain Bank (NBB). All material has been collected from donors from whom a written informed consent for a brain autopsy and the use of the material and clinical information for research purposes had been obtained. Demographic information is displayed in Supplementary Table 11.

#### Autoradiography (ARG)

*In vitro* autoradiography for mHTT was performed using the tritiated ligands [^3^H]CHDI-180 with molar activity (MA) between 70 and 84 Ci/mmol (Pharmaron, UK) or MA = 80 Ci/mmol (Novandi Chemistry AB, Sweden). *In vitro* autoradiography studies were also performed on the same animals of the *in vivo* PET studies using the tritiated version of the ligands (namely [^3^H]CHDI-180, [^3^H]SCH23390, and [^3^H]Raclopride) adapting the previously described procedures^8, 11^. [^3^H]CHDI-180 (MA = 84 Ci/mmol, Pharmaron, UK). [^3^H]SCH23390 (MA = 81 Ci/mmol) and [^3^H]Raclopride (MA = 76 Ci/mmol) were purchased from Novandi Chemistry AB (Sweden).

Twenty *µ*m-thick sections were prepared from brains of hom, het, and age-matched wt mice by using a cryostat, mounted on superfrost slides, and stored at −80°C for a maximum of 2 weeks. On the day of the experiment, slides were adapted to room temperature for 30 min and then equilibrated by immersion into assay buffer (50 mM Tris-HCl pH 7.4; 120 mM NaCl; 5 mM KCl; 2 mM CaCl_2_; 1 mM MgCl_2_) for 20 min at room temperature.

Radioligand solutions for total binding (TB) (0.5 nM (mouse) or 1 nM (human) for [^3^H]CHDI-180, 1 nM for [^3^H]SCH23390, or 2 nM for [^3^H]Raclopride) and non-specific binding (NSB) (0.5 nM of + 1 *µ*M unlabeled compound; 1 or 2 nM + 10 *µ*M unlabeled compound) were prepared in assay buffer. Optimal radioligand concentrations were determined in advance based on the signal-to-background ratio obtained in mouse or human brain tissue. Sections were incubated by immersion into assay buffer containing either only radioligand (TB) or radioligand plus the excess concentration of unlabeled compound (NSB) for 60 min at room temperature. Afterwards, slides were washed three times for 10 min with ice-cold washing buffer (50 mM Tris-HCl, pH 7.4) at 4°C and dipped for three seconds in ice-cold distilled water to remove buffer salts. The slides were dried for 2-3 hours at 30°C and exposed to a Tritium Phosphor Screen (GE Healthcare, Fuji BAS-TR 2025 E) together with calibrated tritium standards (American Radiolabeled Chemicals, ART 0123C, and ART 0123B). Slides and commercial tritium activity standard were exposed for 90 hours ([^3^H]SCH23390 and [^3^H]Raclopride), 96 hours ([^3^H]CHDI-180 at Evotec), or 120 hours ([^3^H]CHDI-180 at MICA). Stored radiation energy on the screen was scanned using a Phosphorimager (GE Healthcare, Typhoon FLA 7000). Densitometric data analysis of radioligand binding was performed using the MCID Analysis 7.1 software (Interfocus Imaging Ltd.) (Evotec) or ImageJ (National Institute of Health, USA) (MICA). Quantification was performed by converting the mean grey values into binding density (fmol/mg) calculated using commercial microscale tritium standards

#### Immunohistochemistry (IHC) in HdhQ80 mouse brains

Twenty-five *µ*m-thick sections were prepared from brains of hom, het, and wt mice by using a cryostat, mounted on 12-well plates (Greiner, Cat. # 665180) and stored at −80°C until further use. The sections were washed three times with DPBS and incubated in 2% H_2_O_2_ for 45 min at room temperature, followed by three times washing with TBS. Antigen retrieval was done by incubation in citrate buffer pH 6.0 for 30 min at 80°C. The sections were permeabilized with 0.1% Triton-X-100, followed by 10% Mouse-to-Mouse Blocking reagent (SkyTek Laboratories, Cat. # MTM015) in 10% normal goat serum and 0.1% Triton-X-100 in TBS for 45 min at room temperature. Sections were incubated overnight at 4°C with mEM48 (1:500, Merck Millipore, Cat. # MAB5374) in 1% normal goat serum and 0.1% Triton-X-100 in TBS. After washing in 0.1% Triton-X-100 in TBS, the sections were incubated with the secondary Goat-Anti-Mouse HRP antibody (1:1000, Abcam, Cat. # ab205719) for two hours at room temperature, followed by washing in 0.1% Triton-X-100 in TBS. The sections were incubated for five min with amplification diluent (0.003% H_2_O_2_ in 0.1 M borate buffer pH 8.5), followed by Biotinyl-Tyramide® amplification kit according to the manufacturer instruction (Perkin Elmer, Cat. # FP1019). After washing in 0.1% Triton-X-100 in TBS the sections were incubated for one hour at room temperature with Streptavidin Dy Light Alexa antibody (1:500; Vector Labs, Cat. # SA-5488) in 0.1% Triton-X-100 in TBS. After washing the sections were incubated overnight at 4°C with anti-NeuN antibody (1:1000, Merck Millipore, Cat. # ABN90P) in 1% normal goat serum and 0.1% Triton-X-100 in TBS. After intense washing, the sections were incubated for two hours at room temperature with the secondary Goat-anti-Guinea Pig antibody in 0.1% Triton-X-100 in TBS. After washing in 0.1% Triton-X-100 in TBS the sections were incubated for 10 min at room temperature with 5 mM DAPI (1:10000, Sigma Aldrich, Cat. # D9542-5MG) in 0.1% Triton-X-100 in TBS. After a brief washing, the sections were allowed to dry and then covered with 400 *µ*l Fluoshield mounting medium (Abcam, Cat. # ab104135), and stored at 4°C.

Automated image acquisition was conducted using the Opera® High Content Screening system and Opera software 2.0.1 (PerkinElmer Inc.), using a 40x water immersion objective (Olympus, NA 1.15, lateral resolution: 0.32 *µ*m/pixel). Image analysis scripts for single-cell analysis were developed using Acapella® Studio 5.1 (PerkinElmer Inc.) and the integrated Acapella® batch analysis as part of the Columbus® system.

#### High-Resolution ARG: Co-registration of [^3^H]CHDI-180 binding and mEM48 staining

Ten *µ*m-thick sections were prepared from PBS-perfused brains of hom, het, and wt mice or human *post-mortem* brain samples by using a cryostat, mounted on superfrost slides, and stored at −80°C for a maximum of 2 weeks. On the day of the experiment, slides were adapted to room temperature for 30 min. The sections were post-fixed with 4% paraformaldehyde for 10 min followed by two times washing with TBS (20 mM Tris-HCl, pH 7.4, 150 mM NaCl). Epitope retrieval was done with 1% formic acid for 10 min followed by washing with TBS.

Slides were equilibrated by immersion into autoradiography assay buffer (50 mM Tris-HCl pH 7.4; 120 mM NaCl; 5 mM KCl; 2 mM CaCl_2_; 1 mM MgCl_2_) for 20 min at room temperature. Radioligand solutions (3 nM ± 10 *µ*M unlabeled compound) were prepared in assay buffer. Optimal radioligand concentrations were determined in advance based on the signal-to-background ratio obtained in mouse or human brain tissue. The solutions were mixed in a Coplin jar (Sigma, S5516-6EA) by gently shaking at 175 rpm for 10 min at room temperature. Sections were incubated by immersion into assay buffer containing either only radioligand (TB) or radioligand plus the excess concentration of unlabeled compound (NSB) for 60 min at room temperature. Afterwards, slides were washed three times for 10 min with ice-cold washing buffer (50 mM Tris-HCl, pH 7.4) at 4°C.

Afterwards, the slides were washed with permeabilization buffer (20 mM Tris-HCl (pH 7.4), 150 mM NaCl, 0.1% TritonX-100) followed by a brief rinse with TBS. Non-specific binding sites were blocked for 1 hour with Mouse on Mouse (M.O.M.) Blocking reagent (Vector Laboratories; MKB-2213) followed by washing with TBS. An additional protein blocking step was done for 20 min with 2.5% horse serum (Fitzgerald; 88R-1020). Sections were incubated for 1 hour with mEM48 (1:500, Merck Millipore, MAB5374) antibody at room temperature in TBS containing 0.1% TritonX-100 (Sigma Aldrich; T9284) and 1% horse serum followed by a washing step in TBS. The ImmPress-AP anti-mouse IgG polymer detection kit (Vector Laboratories; MP-5402) and Vector blue alkaline phosphatase substrate kit (Vector Laboratories; SK-5300) were used as the detection system according to the manufacturer’s instructions. Sections were treated for 10 min with Nuclear fast red (Vectorstain; H3403) for nuclear counterstain followed by rinsing in distilled water.

The slides were allowed to dry and were then covered with NTB emulsion (Kodak/Carestream; 8895666). After drying overnight, the slides were exposed for three weeks at 4°C under lightproof conditions. Photographic development was done by applying the developer X-tol (Kodak; KODAK008) and Vario Fix Powder (Tetenal; S32138) according to the manufactureŕs instructions. Slides were washed with distilled water, allowed to dry for a minimum of 5 hours, and covered in Poly-Mount Mounting Media (Polysciences Europe; 08381-120). Image analysis was done using the PreciPoint M8-S microscope with a 40x air objective.

#### Immunostaining for Imaging Studies

The following primary antibodies were used for immunostaining: monoclonal mouse anti-mHTT (1:100; mEM48 Millipore; Cat. # MAB5374), monoclonal mouse anti-huntingtin (1:1000; 2B4; Millipore, Cat. # MAB5492), polyclonal rabbit anti-GFAP (1:1000; Cat. # Z0334, Dako, Agilent), polyclonal rabbit anti-IBA1 (1:500; Cat. # 019-19741, Wako Chemicals), polyclonal rabbit anti-ZNF-10 (1:300; Cat. # LS-C374589, Lifespan Biosciences), monoclonal rabbit anti-PDE10a (1:2000; EPR22383; Cat. # ab227829; Abcam).

Visualization of mHTT accumulation was performed with two distinct antibodies to explore distinct mHTT species such as inclusion bodies, diffuse nuclear signals, and small nuclear puncta as each antibody may have a higher affinity towards specific species^43^. To assess neuroinflammation/glial reactivity following the intra-striatal injection during the therapy studies, GFAP and IBA1 IHC were executed. For investigating the striatal distribution and efficacy of the ZFP therapy, a co-staining with mEM48 and ZNF-10 was performed. Visualization of the striatal PDE10a levels following ZFP therapy was accomplished by PDE10a staining using the EPR22383 antibody.

Sections were air-dried for 5 min, incubated with 4% paraformaldehyde (PFA) for 10 min for tissue post-fixation, and washed using phosphate-buffered saline (PBS). Next antigen retrieval was performed by placing the slides in a container with citrate buffer into a water bath at 80 °C for 30 min. Then, the slides were cooled at room temperature for 20 min. After rinsing steps with PBS, endogenous peroxidases were inactivated by a 3% H_2_O_2_ solution (for colorimetric IHC). Non-specific binding sites were blocked using 5% normal goat serum (NGS) and 0.5% Triton X-100 in PBS for 30 min and goat anti-mouse Fab fragment IgG (26 *µ*g/ml) for 1 hour (for mEM48 or 2B4); 10% NGS and 0.2% Triton X-100 (GFAP, IBA1), 10% normal donkey serum (NDS) and 0.2% Triton X-100 (PDE10a) in PBS for 1 hour; or 10% NDS and 0.5% Triton X-100 (ZNF10/mEM48) in PBS for 1 hour followed by donkey anti-mouse Fab fragment IgG (1:50; 715-006-151; Jackson Immunoresearch) for 1 hour. Next, sections were incubated overnight at room temperature with the primary antibody in PBS: mEM48 with 3% bovine serum albumin (BSA); 2B4 with 0.1% Triton X-100; polyclonal primary antibodies anti-GFAP or anti-IBA1 with 5% NGS; mEM48 and anti-ZNF-10 with 1% NDS and 1% Triton X-100; anti-PDE10a with 5% NDS and 0.1% Triton X-100. The next day, sections were rinsed with PBS before a 1-hour incubation with the secondary antibody in PBS: horseradish peroxidase (HRP)-conjugated goat anti-mouse (Jackson Immunoresearch, UK) either at 1:500 with 1% NGS for mEM48 staining or at 1:1000 for 2B4 staining; goat anti-rabbit HRP-conjugated (1:500; 111-035-006; Jackson ImmunoResearch) for GFAP or IBA1. Donkey anti-mouse (1:500; AF488-conjugated) and donkey anti-rabbit (1:200; Cy3-conjugated; Jackson Immunoresearch) for ZNF10/mEM48 staining. Donkey anti-rabbit (1:1000; Cy3-conjugated; Jackson Immunoresearch) for PDE10a. Finally, to visualize the binding, sections were either exposed to the colorimetric diaminobenzidine reaction (DAB reagent, Dako) for 10 min and stopped with distilled water for 1 min, dehydrated and mounted with DPX mounting medium (Sigma), or mounted with a solution containing DAPI (4’,6-diamidin-2-fenilindolo) and coverslipped.

For all the markers investigated, whole slices images were acquired at 20X magnification with a whole-slide scanner (Mirax, Zeiss, Germany) and processed with the Pannoramic Viewer (3DHISTECH Ltd, Hungary) or ZEN lite (Zeiss, Germany). Representative images at 100X magnification were obtained with a brightfield microscope (Olympus, Japan) using Olympus CellSens software. Quantitative analyses were performed using Fiji - ImageJ (National Institute of Health, USA) by an experienced investigator blinded to condition after converting the images into grayscale (8-bit) and apply an intensity threshold to remove the background signal.

mHTT aggregate size was determined at different disease stages (3, 6, 9, and 13 months) using mEM48 and 2B4. The area of aggregates in the field of view of different regions-of-interest (ROIs) (namely caudate-putamen, motor cortex, hippocampus, and spinal cord) was measured, and the average was used for statistical analysis. For GFAP and IBA1 immunoreactivity, ROIs (left and right caudate-putamen, CP) were manually drawn on each image and the percentage of the positive area remained after thresholding was measured. For PDE10a immunostaining, ROIs (left and right CP) were manually drawn on each image, the intensity of the signal was measured and compared to the contralateral hemisphere. Similarly, mHTT aggregates and ZFP distribution were investigated in CP bilaterally to assess the efficacy of the treatment in relation to the *in vivo* PET measurements.

### *In vivo* microPET imaging studies

#### Animals

For the naïve zQ175DN experiments, adult male heterozygous (het) and age-matched wt littermates were delivered from Jackson Laboratories (Maine, USA) to MICA (Antwerp, Belgium). For the therapeutic intervention imaging studies, adult male wt and het zQ175DN were received at MICA (Antwerp, Belgium) from Evotec (Hamburg, Germany), blinded for the intervention the animals had undergone at Evotec (Hamburg, Germany). For the LacQ140^I^(*) study, mice have a Q140 m*Htt* knock-in allele which is regulated by Lac operon transcriptional elements, and the mice are also het for the Lac regulator repressor transgene under the β-actin promoter. Binding of the Lac operon and repressor results in whole-body repression of Q140 m*Htt*. The addition of isopropyl β-D-1-thiogalactoside (IPTG) in drinking water de-represses the Q140 allele, allowing full expression of m*Htt*. All mice received 10mM IPTG in their drinking water starting at embryonic day 5 and continuing for the duration of the experiment, or until 2 or 8 months of age [LacQ140^I^(2M) or LacQ140^I^(8M)]. Adult male LacQ140^I^(*)140Q (CHDI-81008005) wt and het mice were shipped from Psychogenics (NJ, USA) to MICA (Antwerp, Belgium) at 12M of age, blinded for genotype and condition.

Animals were single-housed in Eurostandard Type II long cages (Evotec) and individually ventilated cages (MICA) under a 12 hour light/dark cycle in a temperature- and humidity-controlled environment (21 ± 1°C and 55 ± 10% relative humidity) with food and water *ad libitum*. The animals were acclimatized to the facility for at least one week before the start of the procedures. All experiments were conducted during the light phase of the day. For the LacQ140^I^(*)mice requiring IPTG, IPTG was dissolved in the drinking water (2.4 mg/ml) and changed with fresh IPTG water every 3 days. All experimenters at MICA were blinded to treatment allocation, with the group allocations disclosed only upon termination of the analyses.

Animal handling was carried out in accordance with the regulations of the German animal welfare act and the EU legislation (EU directive 2010/63/EU). The study protocol was approved by the local Ethics committee of the Authority for Health and Consumer Protection of the city and state of Hamburg (“*Behörde für Gesundheit und Verbraucherschutz”* BGV, Hamburg) as well as by the Ethical Committee for Animal Testing (ECD #2016-76 and ECD #2018-82) at the University of Antwerp (Belgium).

#### AAV vector construction and production

For the therapeutic intervention studies, ZFP30645flag, targeting specifically the CAG repeat domain, was obtained from Sangamo (ZFP30645flag was termed ZFP-D in original publication^11^) and subcloned via EcoRI/ HindIII into in the adeno-associated virus (AAV) vector pAAV-6P-SWB^44^ under the control of the human synapsin1 promoter (p_hSyn1_). As a control, an inactive ZFP control construct was used; this construct has the deletion of the ZFP DNA binding domain was deleted and only a flag tagged repressor domain (ZFP-ΔDBD) was expressed^11^. Recombinant AAV2/1+2 particles were produced in HEK293 cells co-transfected with the AAV vector carrying the transgene and plasmids containing helper, rep, and cap genes (pDP1rs and pDP2rs, Plasmid Factory). Cells were lysed 48 hours post-transfection, AAV particles released by three freeze-thaw cycles, and purified by iodixanol density centrifugation and a heparin affinity column. Final AAV particles were dialyzed with the AAV Storage buffer (10 mM phosphate buffer+ 180 mM NaCl + 0.001 % Pluronic-F68), aliquoted and stored at −80°C. AAV titers were quantified by qPCR, purity was analyzed by Sypro Ruby gel, and endotoxin levels measured by an EndoZyme® II Recombinant Factor C Endotoxin Detection Assay. Prior to *in vivo* application, the ZFP-expressing AAV vectors were tested *in vitro* in primary cortico-striatal neurons from zQ175DN mice for functionality, i.e. selective m*Htt* downregulation. Moreover, all AAVs were tested *in vivo* for AAV distribution in the striatum after 1 week following intrastriatal injection and the number of microglia and GFAP-positive astroglia quantified in the striatum by IHC in order to evaluate the quality of each AAV batch used for the actual studies.

#### AAV-ZFP *in vivo* striatal injections in mice

Two groups of each 32 zQ175DN male mice at 2 months of age and two groups of each 54 zQ175DN male mice at 5 months of age received bilateral intra-striatal injections (see Fig. 3 and 4 for a graphical representation). Group 1 received the vehicle in the left hemisphere and control ZFP-ΔDBD in the right hemisphere; Group 2 was administered allele-selective ZFP30645 in the left hemisphere and control ZFP-ΔDBD in the right hemisphere. Additionally, one group of 32 wt male mice at 2 months of age and one group of 54 wt male mice at 5 months of age received two intra-striatal injections, one of vehicle in the left hemisphere and another of ZFP-ΔDBD in the right hemisphere. Moreover, two control mice (zQ175DN male animals) from each of the 9 cohorts were injected with allele-selective ZFP30645 in the left and ZFP-ΔDBD in the right hemisphere and analyzed for AAV distribution by IHC 1-week post-injection.

Mice were treated with analgesic and individually anaesthetized with isoflurane and underwent stereotactic surgery (Kopf, Model No. 940). Anesthesia was maintained throughout the surgical procedure. In short, the brain was exposed by drilling a small hole with an electrical drill (Foredom; Model No. H.30) followed by an injection of 4 *µ*l per hemisphere (in total 8X10^9^ GCs; 1 mm anterior, 2.31 mm lateral on right, and 3.6 mm deep (with an angle of 5°) from bregma with flat skull nosebar setting) by using a Hamilton gas tight syringe (model 1801 RN; Cat. No. 7659-0, customized gauge 26 needles) connected to an automated microinjection pump at a constant flow rate of 500 nL/min. Post-injection, the wound was closed and animals were allowed to recover on a heating pad before returning to their holding. Following the appropriate time to recover, animals were shipped to MICA for imaging studies.

#### MRI measurements

Individual magnetic resonance (MR) images of a subset of animals (n = 6 per condition) were acquired at each time point of the therapy studies (3, 6, or 10 months) in zQ175DN mice, as well as at 13 months of age in the LacQ140^I^(*) mice to generate genotype and age dedicated templates for volume-of-interest (VOI) delineation and co-registration purpose as previously described^34^. MRI measurements were performed using a 4.7T (MR Solutions) scanner. Animals were anesthetized using 5% isoflurane (in O_2_/N_2_ 30/70 mixture) and maintained at 1.3-2% of isoflurane (in O_2_/N_2_ 30/70 mixture). Animals were fixed in an MR-compatible holder to immobilize the mouse during imaging and placed in a prone position on the scanner. Body temperature was maintained at 37 ± 1°C utilizing a feedback-controlled warm air circuitry (MR-compatible Small Animal Heating System, SA Instruments, Inc. USA). Three-dimensional (3D) images were acquired with repetition time 2000 ms, echo time 75 ms, and matrix size 256 x 256 x 128. Field of view (FOV) was 20 x 20 x 24 mm^3^ and resolution of 0.0781 x 0.0781 x 0.1875 mm^3^. Data were acquired using ParaVision 6.1 (Bruker, Germany).

#### Radioligand synthesis

##### [^11^C]CHDI-180R – mHTT aggregate detection

The radioligand was prepared using an automated synthesis module (Carbonsynthon I, Comecer, The Netherlands) adapting the previously described procedure^8^ to our system. [^11^C]CHDI-180R was prepared via single-step carbon-11 labeling starting with 0.5 mg of precursor in 0.5 ml of dimethylformamide (DMF), which was reacted with [^11^C]MeI, in the presence of Cs_2_CO_3_ (2.5-3 mg) for 3.5 min at room temperature. To terminate the reaction and to ensure good retention of the compound of interest on the HPLC column, 0.7 ml of water for injection (WFI) was introduced to the reaction mixture, and the resulting crude product was purified on HPLC using a Waters XBridge C18 5*µ*m, 10 mm × 150 mm column (Waters, Belgium), eluted with ethanol/0.05 M sodium acetate pH 5.5 (38/62 V/V) as mobile phase, at a flow rate of 4 ml/min. The collected fraction was sterile-filtered through a 0.22 *µ*m Nalgene PES filter and diluted with sterile saline to decrease the ethanol concentration below 10%. The radiochemical purity was greater than 99% as determined using a Kinetex EVO C18, 5*µ*m, 150 × 4.6 mm (Phenomenex, USA) HPLC column, with acetonitrile/0.05 M sodium acetate pH 5.5 (30/70 V/V) as mobile phase, at a flow of 1.5 ml/min, with UV absorbance set at 280 nm.

##### [^18^F]MNI-659 – PDE10a detection

Radioligand was prepared by adapting our previously described procedure^34^ to the automated AllinOne synthesis module (Trasis, Belgium) using a cassette system built in-house. Synthesis of [^18^F]MNI-659 was accomplished by reacting dried [^18^F]fluoride with the MNI-659 tosylate precursor (7 mg) in dimethyl sulfoxide (DMSO) (1 ml) for 10 min at 90°C. After quenching the reaction mixture with 4 ml of (WFI), [^18^F]MNI-659 was purified by means of semi-preparative HPLC (Waters XBridge C18 5*µ*m, 10 mm × 150 mm column (Waters, Belgium), eluted with acetonitrile/0.1 M ammonium formate 55/45 V/V as mobile phase at a flow rate of 4 ml/min). The collected fraction was diluted with WFI, mixed, and then loaded on a C18 SPE cartridge (Waters, Belgium). After washing the cartridge with 5 ml of WFI, [^18^F]MNI-659 was eluted with ethanol towards the product vial and finally diluted to < 10 % ethanol concentration with sterile saline. Sterile filtration was performed with an in-line 0.22 *µ*m Cathivex GV filter (Merck, Belgium). Final radiochemical purity was > 99 %, as determined using a Waters Xbridge C18 5*µ*m 4.6 mm × 150 mm column (Waters, Belgium) with acetonitrile/0.05 M sodium acetate pH 5.5 (55/45 V/V) as mobile phase, at a flow of 1 ml/min, with UV absorbance set at 230 nm.

##### [^11^C]SCH23390 – D_1_R detection

[^11^C]SCH23390 synthesis was performed on an automated synthesis module (Carbosynthon I, Comecer, The Netherlands) based on the one-pot strategy ^45^ via common *N*-methylation of the desmethyl precursor as previously described ^46^. Briefly, [^11^C]MeI was added to a precooled (−20°C) reaction vessel containing *N*-desmethyl-SCH23390 (1.0 mg) and aqueous NaOH (1 M, 5 *µ*l) in anhydrous DMF/DMSO (ratio 50/50, 300 μL) at room temperature. Following a 3-min reaction at 50°C, the [^11^C]SCH23390 reaction mixture was quenched with 0.9 ml WFI and purified by HPLC (Luna C18(2) 10*µ*m, 10 x 250 mm (Phenomenex, USA); mobile phase: ethanol/0.05 M sodium acetate pH 5.5 50/50 V/V; flow: 3 ml/min). The collected fraction was sterile-filtered through a 0.22 *µ*m Nalgene PES filter and diluted with sterile saline to decrease the ethanol concentration below 10 %. The radiochemical purity was greater than 99 %, as determined using a Luna C18 5*µ*m 4.6x 150 mm (Phenomenex, USA) HPLC column, with acetonitrile/0.05 M sodium acetate pH 5.5 (30/70 V/V) as mobile phase, at a flow of 1 ml/min, with UV absorbance set at 280 nm.

##### [^11^C]Raclopride – D_2/3_R detection

Radioligand was prepared using an automated synthesis module (Carbonsynthon I, Comecer, The Netherlands) adapting the previously described procedure^47^ to our system. Briefly, [^11^C]Raclopride was synthesized via common *O*-methylation of the phenolic hydroxyl moiety. [^11^C]Methyl triflate was trapped in a V-shaped reaction vial containing *O*-desmethyl-raclopride (1 mg) and aqueous NaOH (1 M, 5 *µ*l) in reagent grade acetone (300 *µ*l) at room temperature. Ninety seconds after the end of trapping, the reaction mixture was quenched with water for injection (600 *µ*l) and purified using isocratic semi-preparative reverse phase HPLC (Waters XBridge™ Prep C18 5 *µ*m 10×150 mm, λ = 254 nm; mobile phase: ethanol/phosphate buffer pH 5.5 40/60 V/V, 3 ml/min). The collected fraction was sterile-filtered through a 0.22 *µ*m Nalgene PES filter and diluted with sterile saline to decrease the ethanol concentration below 10 %. Radiochemical purity (RCP) was > 99% as determined by analytical isocratic reverse-phase HPLC (Waters Symmetry^®^ C18 3.5 *µ*m 4.6×50 mm, λ = 220 nm; mobile phase: sodium 1-heptanesulfonate 2 g/L pH 3.9/acetonitrile 70/30 v/v, 1 ml/min).

#### Radiometabolite analysis for [^11^C]CHDI-180R

To evaluate the *in vivo* metabolism of [^11^C]CHDI-180R, a radiometabolite analysis was performed on a young (4-month-old) and an aged (10-month-old) cohort of wt and het zQ175DN mice (*n* = 3-4 for each genotype, time point, and age) adapting the previously described procedure^48, 49^. Mice were injected via the lateral tail vein with [^11^C]CHDI-180R (11.7 ± 2.9 MBq in 200 μl) while keeping cold mass within tracers condition (<1.5 *µ*g/kg). Animals were anesthetized, and blood was withdrawn via cardiac puncture and the brain was rapidly removed by dissection at 15, 25, and 40 min p.i. Following centrifugation of blood at 2377 × rcf at 4°C for 5 min, both plasma and brain samples were counted in a gamma counter (Wizard^2^, PerkinElmer) to determine the total radioactivity. Next, equal amounts of ice-cold acetonitrile and 10 *µ*l of cold reference (1 mg/ml) were added to the plasma samples. The brain samples were homogenized in ice-cold acetonitrile (1ml), 10 *µ*l of cold reference compound (CHDI-180) was added as well. After another centrifugation at 2377 × rcf at 4°C for 5-min, the supernatant was separated from the precipitate and both fractions were counted in the gamma counter to calculate the extraction efficiencies in both plasma and brain samples. Extraction efficiency did not differ among genotypes and ages for plasma (4 months: wt = 94.5 ± 0.4%, het = 94.1 ± 0.6%; 10 months: wt = 94.2 ± 1.2%, het = 94.0 ± 0.8%, mean ± s.d.) and brain samples (4 months: wt = 84.8 ± 1.4%, het = 82.2 ± 4.2%; 10 months: wt = 85.6 ± 2.5%, het = 84.2 ± 4.7%, mean ± s.d.). Next, 100 μl of supernatant were loaded onto a pre-conditioned reverse-phase (RP)-HPLC system (Kinetex, 150×4.6 mm, 5 μm HPLC column + Phenomenex security guard pre-column) and eluted using an isocratic system comprised of 0.1% TFA in H_2_O and acetonitrile: NaOAc 0.05M pH 5.5 (80/20 v/v) buffer at a flow rate of 1 ml/min. RP-HPLC fractions were collected at 0.5 min intervals for 8 min and radioactivity was measured in the gamma counter. The radioactivity was expressed as a percentage of the total area of the peaks based on the radiochromatograms. Blood and brain samples spiked *in vitro* with 37 kBq of radiotracer indicated that no degradation occurred during procedural work-up for any sample (parent = 99.7 ± 0.25%).

#### PET imaging

##### Image acquisition

Dynamic microPET/Computed tomography (CT) images were acquired using two virtually identical Siemens Inveon PET/CT scanners (Siemens Preclinical Solution, Knoxville, USA) as previously described^46, 48, 49^. Animals were anesthetized using isoflurane in medical oxygen (induction 5%, maintenance 1.5%) and catheterized in the tail vein for intravenous (i.v.) bolus injection of the tracer. Animals were placed on the scanner bed with the full body in the PET scanner’s field of view (FOV) to allow the extraction of the image-derived input function (IDIF) from the left ventricle as previously described^24, 25^. Bolus injection of radiotracer occurred over a 12-second interval (1 ml/min) using an automated pump (Pump 11 Elite, Harvard Apparatus, USA) at the onset of the dynamic microPET scan. Information regarding molar activity injected radioactivity, injected mass, body weight, and age on scan day for each radioligand at different time points and studies are reported in Supplementary Tables 1, 4, 5-7. Radioligands were injected with activity as high as possible to obtain good image quality while keeping the cold mass as low as possible in order not to violate tracer conditions (namely 1.25 *µ*g/kg for [^11^C]CHDI-180R, 1 *µ*g/kg for [^18^F]MNI-659, 2 *µ*g/kg for [^11^C]SCH23390, 1.5 *µ*g/kg for [^11^C]Raclopride). PET data were acquired in list mode format. Dynamic scans lasted 60 min for [^11^C]CHDI-180R and [^11^C]Raclopride, while a 90 min acquisition was performed for [^18^F]MNI-659 and [^11^C]SCH23390. PET scans were followed by a 10 min 80 kV/500 μA CT scan on the same gantry for attenuation correction and coregistration purposes. Acquired PET data were reconstructed into 33 or 39 (for 60 or 90 min acquisition, respectively) frames of increasing length (12 x 10s, 3 x 20s, 3 x 30s, 3 x 60s, 3 x 150s, and 9 or 15 x 300s) using a list-mode iterative reconstruction with proprietary spatially variant resolution modeling in 8 iterations and 16 subsets of the 3D ordered subset expectation maximization (OSEM 3D) algorithm^50^. Normalization, dead time, and CT-based attenuation corrections were applied. PET image frames were reconstructed on a 128 x 128 x 159 grid with 0.776 x 0.776 x 0.796 mm^3^ voxels.

##### Image processing

Image analysis was performed with PMOD 3.6 software (Pmod Technologies, Zurich, Switzerland) applying a CT-based pipeline for the longitudinal natural history study and an MR-based pipeline for the therapeutic and LacQ140^I^(*) studies. When we applied the CT-based pipeline, spatial normalization of the PET/CT images was done through brain normalization of the CT image to the CT/MRI template with predefined volumes-of-interest (VOIs) adapting the previously described procedure^34^. The spatial transformations were applied to the dynamic PET images and assessed for accuracy following spatial transformation. Using the VOI template adapted from the Waxholm atlas^51^ (as shown in Extended Fig. 3), time-activity curves (TACs) for the striatum, motor cortex, hippocampus, thalamus, and cerebellum were extracted from the dynamic PET images to perform kinetic modeling.

Since we previously observed that the use of magnetic resonance imaging (MRI) templates for spatial normalization and VOI definition improves the accuracy of the regional quantification of PET data with focal uptake, the therapeutic and LacQ140^I^(*) studies were processed using an MR-based pipeline^34^. VOIs were manually adapted from the Waxholm atlas^51^ to match each genotype and age-specific MR template. TACs for the striatum, motor cortex, hippocampus, thalamus, and cerebellum were extracted from the dynamic PET images in order to perform kinetic modeling.

For analysis of spinal cord and peripheral tissue (heart, liver, muscle, and adipose tissue), VOIs were manually drawn on the individual CT images, and TACs were extracted from the dynamic scans for regional quantification.

##### Kinetic modeling

In zQ175DN and Q140 mouse models, mHTT accumulates in all brain structures^8, 13, 21, 40^ and no suitable reference region for relative quantification could be identified. Hence absolute quantification for [^11^C]CHDI-180R was performed to calculate the total volume of distribution based on image-derived input function (*V*_T (IDIF)_) as a non-invasive surrogate of the *V*_T_. Kinetic modeling fitted regional TACs using the Logan model^27^ and the image-derived input function (IDIF) with the start of the linear regression (*t*\*) calculated according to the maximum error criterion of 10%. The IDIF was obtained from the whole blood activity derived from the PET images by generating a region-of-interest (threshold-based 50% of max) in the lumen of the left ventricle as previously described^24, 48^. Since only negligible metabolism of [^11^C]CHDI-180R was observed in different genotypes and ages (parent compound >95%), no correction for radiometabolites was applied.

Parametric *V*_T (IDIF)_ maps were generated through voxel-based graphical analysis (Logan)^27^ using the IDIF as input function, and were then cropped using the brain mask of the MRI template, represented as group averages, and overlaid onto a 3D mouse brain template for anatomical reference. Individual images were smoothed with an isotropic gaussian filter (0.5 mm in full width at half maximum). For the longitudinal natural history study, voxel-based analysis with Statistical Parametric Mapping (SPM) using SPM12 (Wellcome Centre for Human Neuroimaging) was performed on het zQ175DN mice to evaluate the voxel-based changes with disease progression. Data from zQ175DN het mice were compared between time points in order to determine longitudinal changes in [^11^C]CHDI-180R *V*_T (IDIF)_. Statistical *t* maps were generated for a peak voxel threshold of *P*=0.01 (uncorrected) and a cluster threshold of 100 voxels (0.8 mm^3^). Only significant clusters with *P*<0.01 were considered.

For the quantification of [^18^F]MNI-659, [^11^C]SCH23390, and [^11^C]Raclopride the binding potential (*BP*_ND_) was determined by fitting the regional TACs using the simplified reference tissue modeling (SRTM)^52^. The striatum was selected as the receptor-rich region and the cerebellum the receptor-free region (reference region)^34, 46^. Parametric *BP*_ND_ maps were generated using SRTM2^53^ with the *k*_2_’ as calculated with SRTM^52^. The individual images were smoothed with an isotropic gaussian filter (0.5 mm in full width at half maximum), cropped using the brain mask of the MRI template, represented as group averages, and overlaid onto each condition- and age-specific 3D brain template for anatomical reference.

In the early and late ZFP intervention studies, the therapeutic response of each molecular target was estimated as follows according to Fig. 3b and 4b:

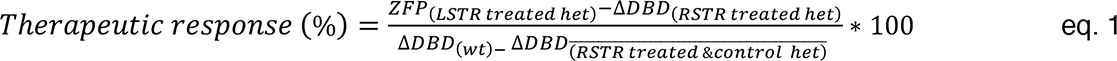

Where LSTR and RSTR represent the left and right striatum, respectively.

In the LacQ140^I^(*) studies, the mHTT lowering response of [^11^C]CHDI-180R was estimated as follows:

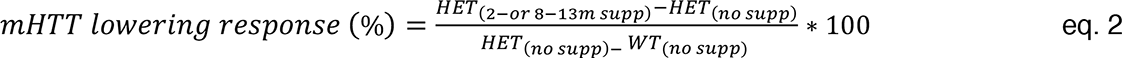

#### Tissue collection

zQ175DN mice were euthanized at 3, 6, 9, 10, 13, and 16 months of age. LacQ140^I^(*) mice were euthanized at 13 months of age and the cerebellum was removed for MSD analysis. For the longitudinal zQ175DN natural history and LacQ140^I^(*) studies, brains, as well as the cervical spinal cord, were quickly removed from the skull and fresh-frozen in 2-metylbuthane at −35°C for 2 min and further preserved at −80°C until use. Cerebral serial coronal sections of 20 *µ*m thickness were collected starting at 1.10 mm (striatum) and −1.46 mm (dorsal hippocampus) from bregma according to the Paxinos and Franklin mouse brain atlas^54^ in triplicate using a cryostat (Leica, Germany) and dry mounted on coated Superfrost Plus slides (Thermo Fischer Scientific, USA). Additionally, for a subset of animals, the cervical spinal cord was also dissected and sections of 30 *µ*m thickness were collected.

For the therapeutic studies, animals from each condition were equally assigned to either fresh-frozen or PFA-perfused groups for post-mortem assessment. For the fresh-frozen group, brains were removed and frozen in isopentane at −35°C for 2 min and subsequently preserved at −80°C. For the PFA-perfused group, deep anesthesia was induced with ketamine/xylazine (120/15 mg/kg). Before perfusion, loss of toe pinch reflex was assessed to ensure that the correct level of anesthesia was achieved. Animals were intracardially perfused first with 12 ml of PBS followed by 4% PFA using an automatic pump (flow rate 120 ml/h). Then, brains were removed and placed into 4% PFA for 2 hours. Next, brains were transferred into 30% sucrose with 0.2% NaN_3_ solution for 72 hours at 4°C, frozen, and subsequently preserved at −80°C. Serial coronal sections of 10 *µ*m thickness were collected (from bregma = 1.70 mm until bregma = −0.22 mm^54^) in triplicate by a cryostat and dry mounted on glass microscope slides. Hematoxylin and eosin (H&E) staining was performed to identify the anatomical region corresponding to the striatal injected area and select the correct slides to be used for *in vitro* assessment.

#### Meso Scale Discovery (MSD) analysis of HTT levels

##### Tissue preparation

Four punches from frozen striata were harvested from coronal frozen sections of 0.5 mm height: two punches with a diameter of 1.5 mm and 0.5 mm height close to the injection site and two punches with a diameter of 1 mm and 0.5 mm height adjacent to the first punches. For LacQ140^I^(*), the cerebellum was used for MSD analysis. The collected striatal punches were lysed in 100 *µ*l of tissue lysis buffer (20 mM Tris (pH 7.5), 150 mM NaCl, 1 mM EDTA, 1 mM EGTA, 1% Triton X-100, 10 mM NaF, 1 mM PMSF, Phosphatase Inhibitor Cocktail II (Sigma), Phosphatase Inhibitor Cocktail III (Sigma), Protease Inhibitors (Roche Diagnostics)) using a FastPrep-24 tissue homogenizer (MP Biomedicals) for up to three times 30 s cycles. Crude lysates were centrifuged three times (10 min at 16,000 rcf and 4°C), and the supernatant was collected and transferred to a new tube after each centrifugation step. Total protein concentration was determined by the bicinchoninic acid assay (BCA; Thermo Scientific) and adjusted to 1 mg/ml using lysis buffer. Homogenates were aliquoted, snap-frozen, and stored at −80°C until analysis.

##### Meso Scale Discovery Analysis

MSD plates (384-well) were coated overnight at 4°C with 10 *µ*l of coating antibody in carbonate-bicarbonate coating buffer (15 mM Na_2_CO_3_/35 mM NaHCO_3_, pH 9.6) per well. Plates were washed 3 times with 35 *µ*l of wash buffer (0.2% Tween-20 in PBS) per well and blocked with 35 *µ*l 2 % probumin/ 0.2 % Tween-20 in PBS for 1 hour at room temperature with rotational shaking. Striatal extracts were diluted to 0.5 mg/ml in a mixture of 50% tissue lysis buffer and 50% blocking buffer. After an additional washing step, 10 *µ*l of per sample were transferred to each well of the antibody-coated MSD plate and incubated with shaking for 1 hour at room temperature. After disposal of samples and four wash cycles, 10 *µ*l of the detection antibody were added to each well and incubated with shaking for 1 hour at room temperature. After three times washing 35 *µ*l of read buffer T with surfactant (Meso Scale Discovery) were added to each well and the plate imaged on a Sector Imager 6000 (Meso Scale Discovery) according to manufacturers’ instructions. The following antibody combinations were used: for soluble mHTT assay (5 *µ*g/ml 2B7 / 5 *µ*g/ml MW1-ST (SULFO-tag)); for detection of aggregated mHTT (4 *µ*g/ml MW8 / 1 *µ*g/ml 4C9-ST) and for detection of total mouse HTT (8 *µ*g/ml CHDI-90002133, mouse-specific polyPro mAb / 1 *µ*g/ml D7F7-ST). Samples were quantified using the following recombinant protein standards: large human fragment HTT-Q73, aa 1-573, FLAG N-term (for assay 2B7 / MW1-ST)^36^, human HTT-Q46, aa 1-97, N-term MBP, C-term 6His, with thrombin cleavage site, thrombin-digested to obtain aggregated HTT (for MW8 / 4C9-ST and MW8)^37^, and mouse HTT-Q7, aa 1-3120, TEV, FLAG C-term (for assay 2133/D7F7).

#### Analysis of AAV distribution

In order to analyze the striatal distribution of AAV2 1+2 vectors, 2-month-old zQ175DN het mice were injected with ZFP constructs containing an HA-tag, which allows for better determination of positive cells by IHC. For perfusions at the age of 4 months, mice were deeply anesthetized by intraperitoneal injection of ketamine/xylazine mixture (120 mg/15 mg per kg in 15 μl/g body weight). Mice were transcardially perfused with 30 ml of ice-cold PBS followed by 50 ml of 4% paraformaldehyde. Brain samples were removed, post-fixed overnight at 4°C, and cryoprotected in 30% sucrose solutions until saturated. Whole brains were embedded in TissueTek and stored at −80°C. Coronal sections of 25 *µ*m were cut using a cryostat, collected as free-floating in 24-well plates. All stainings were performed with free-floating sections. Sections were permeabilized in 0.3% Triton X-100/PBS, blocked in 10% normal goat serum/PBS, and incubated with the primary antibodies diluted in 1% normal goat serum, 0.1% Triton X-100 in PBS at 4°C overnight: Primary antibodies used were anti-NeuN (1:1000, Millipore, ABN90P, lot#3226535), polyclonal rabbit anti-HA (C29F4) (1:400, Cell signaling, 3724S, lot#9). After three washes in PBS sections were incubated with secondary antibodies (anti-Rabbit IgG CF™ 568, 1:1000, Sigma Aldrich, SAB4600084, and Anti-Guinea Pig CF™ 647, 1:1000, Sigma Aldrich, SAB4600180) for 2 hours at room temperature. Subsequently, sections were washed twice in 0.1% Triton X-100/PBS, incubated with DAPI (1:10,000, Sigma Aldrich, D9542), washed once in 0.1% Triton X-100/ PBS, and mounted using 10 mM Tris-buffered saline pH 7.4 in 24-well glass-bottom plates (24 Well SensoPlate^TM^, Greiner, #662892) suitable for imaging with the Opera High Content Screening System (PerkinElmer Inc.). For fluorescence preservation, sections were covered with an aqueous mounting medium (Anti-Fade Fluoroshield Mounting Medium, Abcam, 104135).

Automated image acquisition was conducted using the Opera® High Content Screening system and Opera software 2.0.1 (PerkinElmer Inc.), using a 40x water immersion objective (Olympus, NA 1.15, lateral resolution: 0.32 *µ*m/pixel). Image analysis scripts for HA+/NeuN+ cells were developed using Acapella® Studio 5.1 (PerkinElmer Inc.) and the integrated Acapella® batch analysis as part of the Columbus® system.

### Statistical analysis

Statistical analysis was performed in GraphPad Prism v9.1 (GraphPad Software) and JMP Pro 14 (SAS Institute Inc.). Data are expressed as the mean ± standard deviation (s.d.) unless otherwise indicated in the figure legends. To choose the appropriate statistical test, data were checked for normality using the Shapiro-Wilk test. If the normality test was not passed, non-parametric statistical tests were used. Longitudinal analysis of each PET readout was performed using linear mixed-effects models with each radioligand quantification as the primary endpoint. Genotype, cohort, time point, region, and treatment (when applicable) as fixed factors, while subjects as a random effect. Interaction effects (genotype*time, cohort*time, treatment*time, and treatment*region) were evaluated as well. Comparisons were performed to evaluate regional temporal and genotypic differences as well as treatment effects. Correlation coefficients were calculated with Pearson’s correlation analysis. Sample size calculations at desired therapeutic effects were performed in G*Power software (http://www.gpower.hhu.de/). Statistical significance was set at *P* < 0.05, with the following standard abbreviations used to reference *P* values: ns, not significant; ^#^ *P* < 0.1; * *P* < 0.05; ** *P* < 0.01; *** *P* < 0.001; **** *P* < 0.0001. Detailed statistical information for each experiment is provided in the corresponding figure legends.

## Acknowledgments

The authors thank Philippe Joye, Caroline Berghmans, Eleni Van der Hallen, Romy Raeymakers, Silvia Incardona, and Annemie Van Eetveldt of the Molecular Imaging Center Antwerp (MICA) for their important and valuable technical support, Evelyn Galstian, Maya Bader, and Brenda Lager for project and animal management support and administration, as well as Dr. Simon Noble for help with proofreading the manuscript. DB is supported by a post-doctoral fellowship from the Research Foundation Flanders (FWO, 1229721N). Antwerp University, Belgium supported the work through a partial assistant professor position to JV and a full professor position to SSta.

## Author contributions

D.B., J.B., L.L., M.H., A.G., C.D., J.V., S.St. and I.M.-S. contributed to the conception and design of the studies. F.P., F.H., and S.Sc. contributed to *in vitro* assays. F.H., C.B., P.J., M.P., and M.M. contributed to the chemistry design of ligands and synthetic routes. D.B. and M.H. contributed to *in vitro* studies. M.H. contributed to the development of *in vitro* co-registration studies. A.G. and B.H. contributed to ZFP injection studies. D.B., T.V., and A.V.d.L. contributed to the MR data. S.D.L. contributed to the synthesis of radioligands. D.B., A.M., F.Z., J.V., S.St. contributed to *in vivo* PET studies. D.B. and F.Z. contributed to *post-mortem* studies. D.B., J.B., L.L., A.G., L.M., V.K., Y.W., D.M., M.S., J.V., C.D., S.St., and I.M.-S. supervised the experiments. D.B., J.B., M.H., L.L., A.G., L.M., V.K., Y.W., D.M., M.S., J.V., C.D., S.St. and I.M.-S. contributed to the interpretation of the results. D.B. prepared the Figures. D.B., J.B., M.H., A.G., and I.M.-S. wrote the manuscript. All authors contributed to and have approved the final manuscript.

## Data availability

All requests for data will be promptly reviewed by the institutions involved to verify whether the request is subject to any intellectual property or confidentiality obligations. If deemed necessary, a material transfer agreement between requestor and institutions involved may be required for sharing of some data. Any data that can be freely shared will be released.

## Extended Data

**Extended Data Fig. 1.**
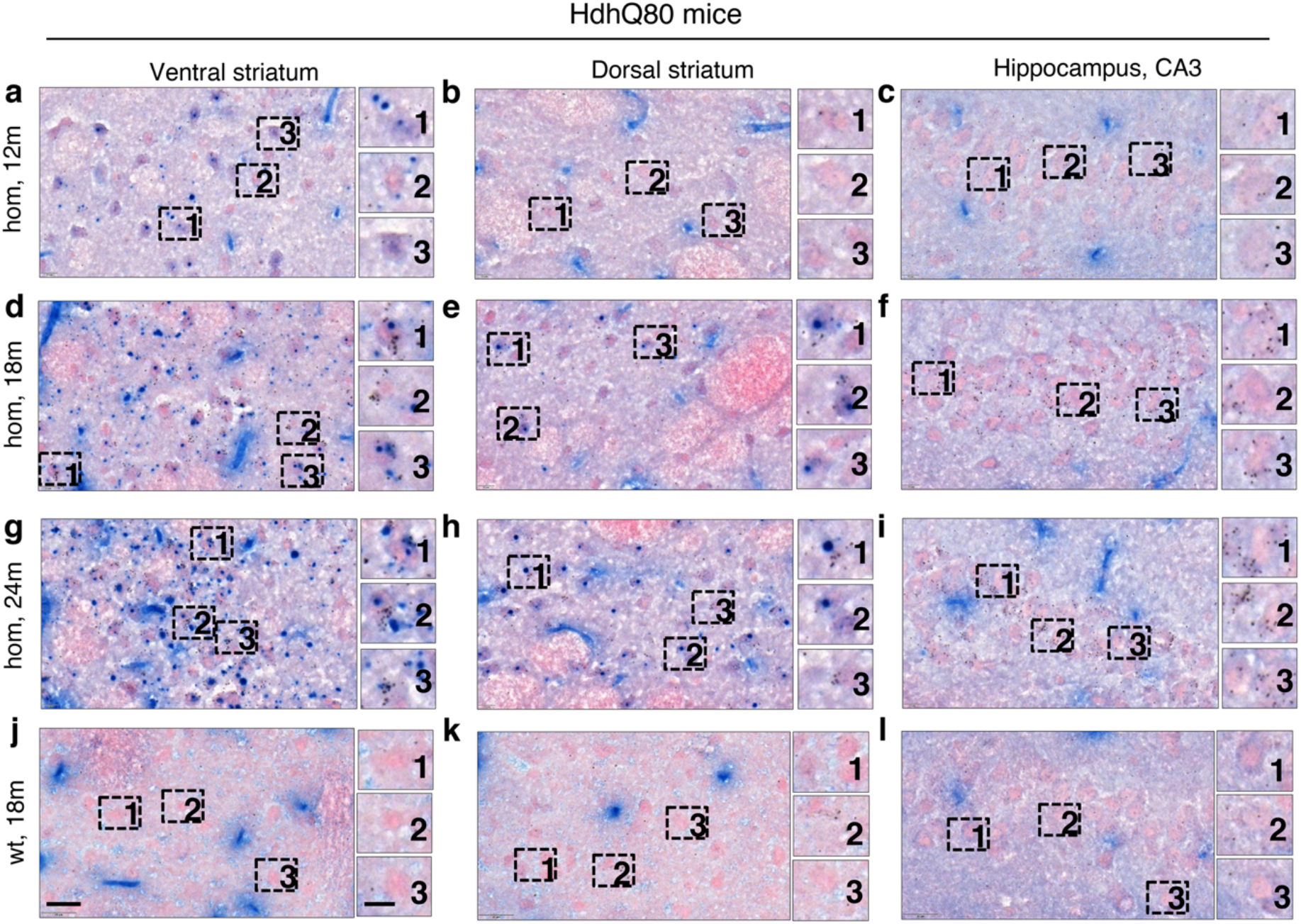
Absence of colocalization of [^3^H]CHDI-180 binding to mHTT inclusions in Hdh80 mice. **a-i**, [³H]CHDI-180 binding and mHTT inclusions (mEM48) in 12-, 18-, and 24-month-old hom HdhQ80 mice in the ventral striatum (**a**, **d**, **g**), dorsal striatum (**b**, **e**, **h**), and hippocampal CA3 (**c**, **f**, **i**). Age-dependent increase in the number of [³H]CHDI-180 silver grain signals that follow the appearance of mEM48-positive mHTT inclusions in dorsal and ventral striatum without co-registering with mHTT inclusions (**a**-**b**, **d**-**e**, **g**-**h**). In hippocampal CA3, no positive mEM48 staining detectable up to 24m; however, an age-dependent increase in the number of [³H]CHDI-180 silver grains is observed (**c**, **f**, **i**). [³H]CHDI-180 binding and mHTT inclusions (mEM48) in 18-month-old wild-type HdhQ80 mice in the ventral striatum (**j**), dorsal striatum (**k**), and hippocampal CA3 (**l**). No [³H]CHDI-180 silver grain signals detectable. [^3^H]CHDI-180 binding, black silver grains; mHTT inclusions (mEM48), blue; background tissue (Nuclear Fast Red), pink. Scale bar, 20 *µ*m; inset, 10 *µ*m. Related to Figure 1.

**Extended Data Fig. 2.**
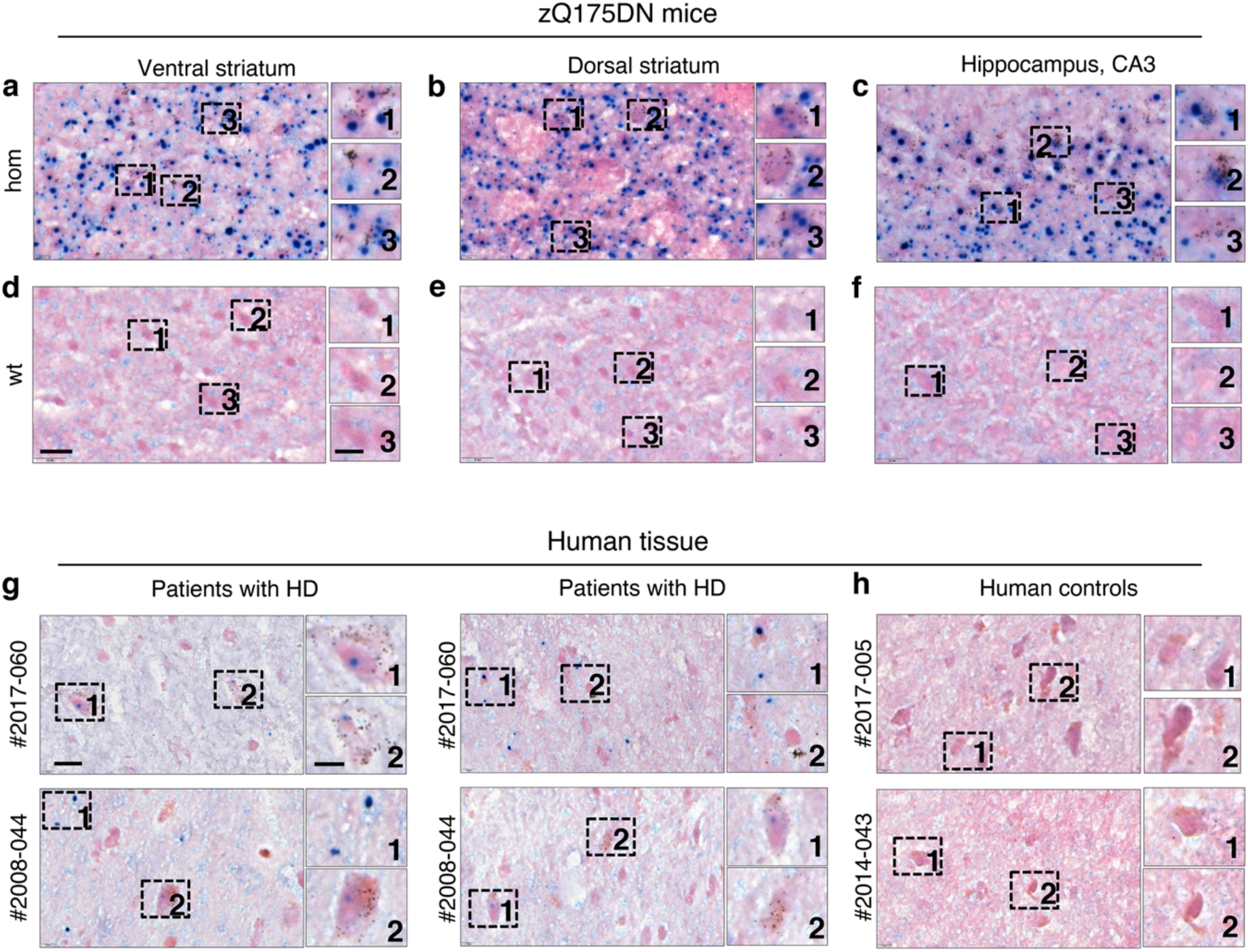
Absence of colocalization of [^3^H]CHDI-180 binding to mHTT inclusions in zQ175DN mice and human tissue. **a-c**, [³H]CHDI-180 binding and mHTT inclusions (mEM48) in 12-month-old hom zQ175DN mice in the ventral striatum (**a**), dorsal striatum (**b**), and hippocampal CA3 (**c**). [³H]CHDI-180 silver grain signal was detectable in close vicinity to mEM48-positive signal but never co-register with mHTT inclusion bodies, although it was partially co-registered with more diffuse appearing mEM48-positive signal. No [³H]CHDI-180 silver grain signals detectable in 12-month-old wt zQ175DN mice (**d**-**f**). **g**, Colocalization of [³H]CHDI-180 binding and mHTT inclusions (mEM48) in the *post-mortem* frontal cortex sections of two patients with HD (#2017-060 and #2008-044) showed similar results as obtained with HD mouse models. [³H]CHDI-180 silver grain signal clustered around cells with mEM48-positive or mEM48-negative signal, but never co-register with mHTT inclusion bodies. **H**, No [³H]CHDI-180 silver grain signals detectable in the *post-mortem* frontal cortex sections of non-demented human controls (#2017-005 and #2017-043). [^3^H]CHDI-180 binding, black silver grains; mHTT inclusions (mEM48), blue; background tissue (Nuclear Fast Red), pink. Scale bar, 20 *µ*m; inset, 10 *µ*m. Related to Figure 1.

**Extended Data Fig. 3.**
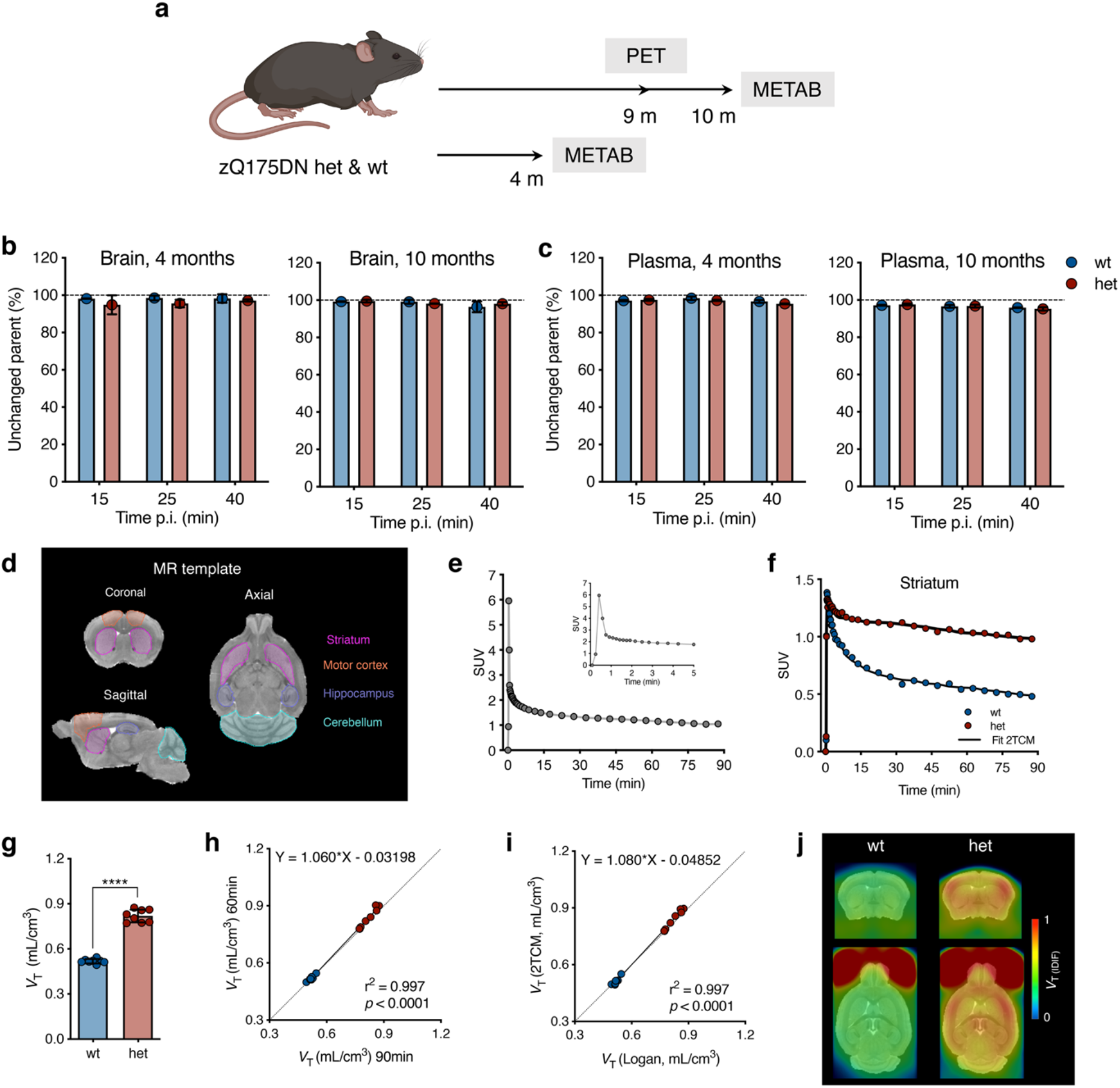
PET imaging properties of [^11^C]CHDI-180R in zQ175DN wt and het mice. **a**, Timeline overview and endpoints in zQ175DN wt and het mice for characterization of [^11^C]CHDI-180R PET imaging. **b**,**c**, Percentage of unchanged parent radioligand detected at 15 (wt, n = 3; het, n = 3), 25 (wt, n = 3; het, n = 3), and 40 (wt, n = 3; het, n = 3) min following i.v. injection into the brain (**b**) and plasma (**c**) in zQ175DN wt and het mice at 4 and 10 months of age. The dotted line represents a completely unchanged radioligand (100%). Data are presented as a percentage of total CPM in the radiochromatogram and shown as mean ± s.d., all points shown. **d**, Volumes of interest applied for quantification to microPET imaging studies. Volumes of interest are co-registered to the MRI template for anatomical reference **e**, Representative [^11^C]CHDI-180R image-derived input function of one zQ175DN het mouse at 9 months of age during a 90-min microPET acquisition. **f**, Representative striatal standardized time-activity curves of zQ175DN wt and het mice at 9 months of age with the curve fitting of the two-tissue compartment model (2TCM; solid lines) over a 90-min microPET scan. **g**, Regional [^11^C]CHDI-180R *V*_T (IDIF)_ quantification in zQ175DN wt and het at 9 months (wt, n = 8; het, n = 8) of age based on 2TCM. Two-tailed unpaired t-test with Welch’s correction; **** *P* < 0.0001. Data are shown as mean ± s.d., all points shown. **h**, Correlation between striatal mHTT binding measured with [^11^C]CHDI-180R microPET during 90 min and 60 min acquisition in zQ175DN wt and het mice at 9 months of age. Two-tailed Pearson correlation analysis; R^2^ = 0.997; *P* < 0.0001. Dotted line represents the identity line. **i**, Correlation between striatal mHTT binding measured with [^11^C]CHDI-180R microPET using 2TCM and Logan plot models for *V*_T (IDIF)_ estimation during 60 min acquisition in zQ175DN wt and het mice at 9 months of age. Two-tailed Pearson correlation analysis; R^2^ = 0.997; *P* < 0.0001. Dotted line represents the identity line. **j**, Mean [^11^C]CHDI-180R *V*_T (IDIF)_ parametric images based on Logan plot of zQ175DN wt and het mice (wt, n = 8; het, n = 8) at 9 months of age. PET images are co-registered to the MRI template for anatomical reference. Coronal (top) and axial (bottom) planes are shown. Related to Figure 2.

**Extended Data Fig. 4.**
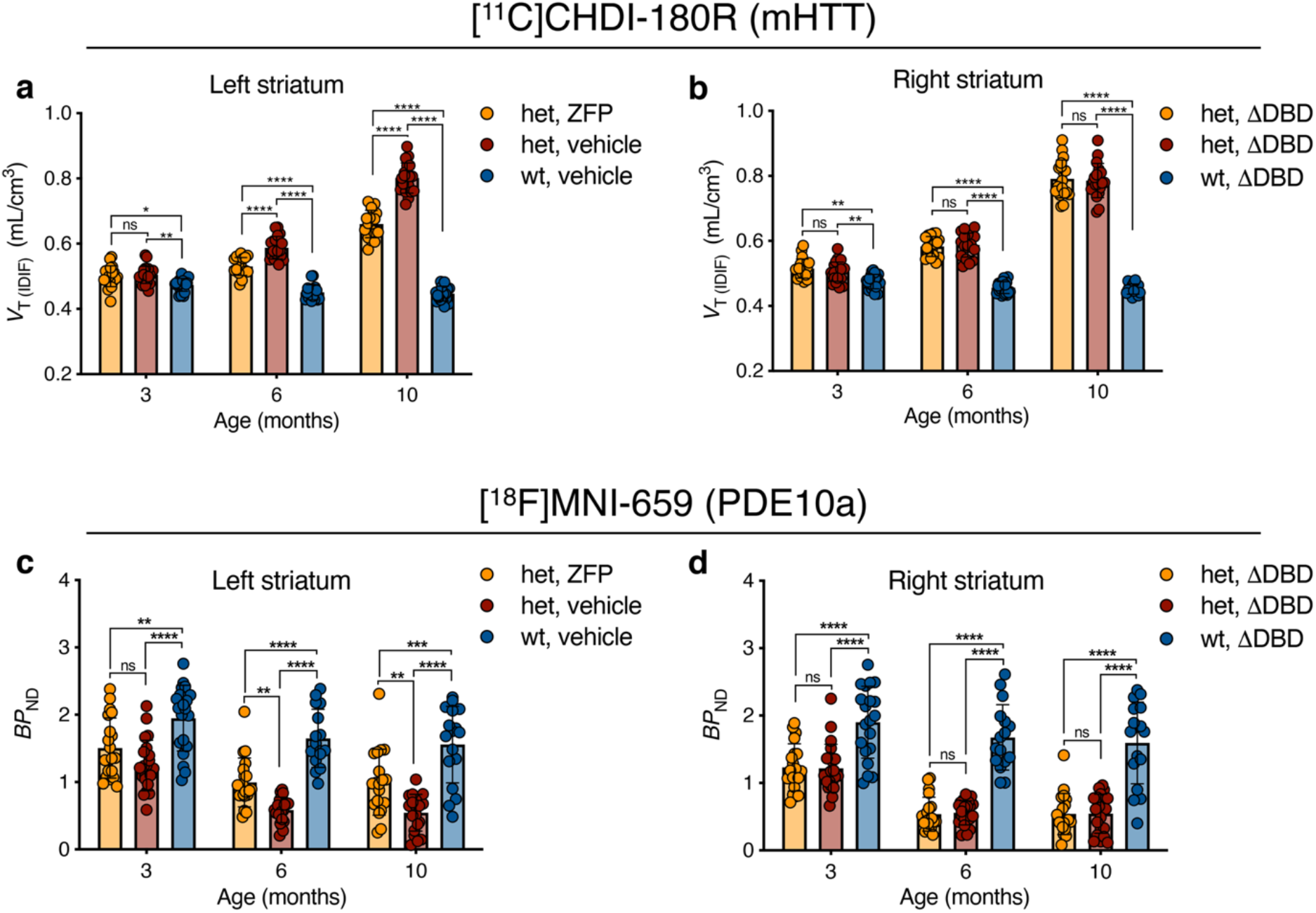
Longitudinal striatal PET imaging quantification following early ZFP intervention in zQ175DN mice. **a**,**b**, Quantification of [^11^C]CHDI-180R *V*_T (IDIF)_ (mHTT inclusions) in left (**a**) and right (**b**) striatal hemispheres of zQ175DN wt vehicle, het vehicle, and het ZFP-treated mice at 3 months (het ZFP, n = 21; het vehicle, n = 21; wt vehicle, n = 20), 6 months (het ZFP, n = 18; het vehicle, n = 19; wt vehicle, n = 19), and 10 months (het ZFP, n = 18; het vehicle, n = 19; wt vehicle, n = 19) of age following striatal injection at 2 months of age. Repeated measures with linear mixed model analysis with Tukey-Kramer correction; * *P* < 0.05, ** *P* < 0.01, **** *P* < 0.0001. Data are shown as mean ± s.d., all points shown. **c**,**d**, Quantification of [^18^F]MNI-659 *BP*_ND_ (PDE10a) in left (**c**) and right (**d**) striatal hemispheres of zQ175DN wt vehicle, het vehicle, and het ZFP-treated mice at 3 months (het ZFP, n = 21; het vehicle, n = 22; wt vehicle, n = 20), 6 months (het ZFP, n = 21; het vehicle, n = 21; wt vehicle, n = 18), and 10 months (het ZFP, n = 20; het vehicle, n = 20; wt vehicle, n = 18) of age following striatal injection at 2 months of age. Repeated measures with linear mixed model analysis with Tukey-Kramer correction; ** *P* < 0.01, **** *P* < 0.0001. Data are shown as mean ± s.d., all points shown. Related to Figure 3.

**Extended Data Fig. 5.**
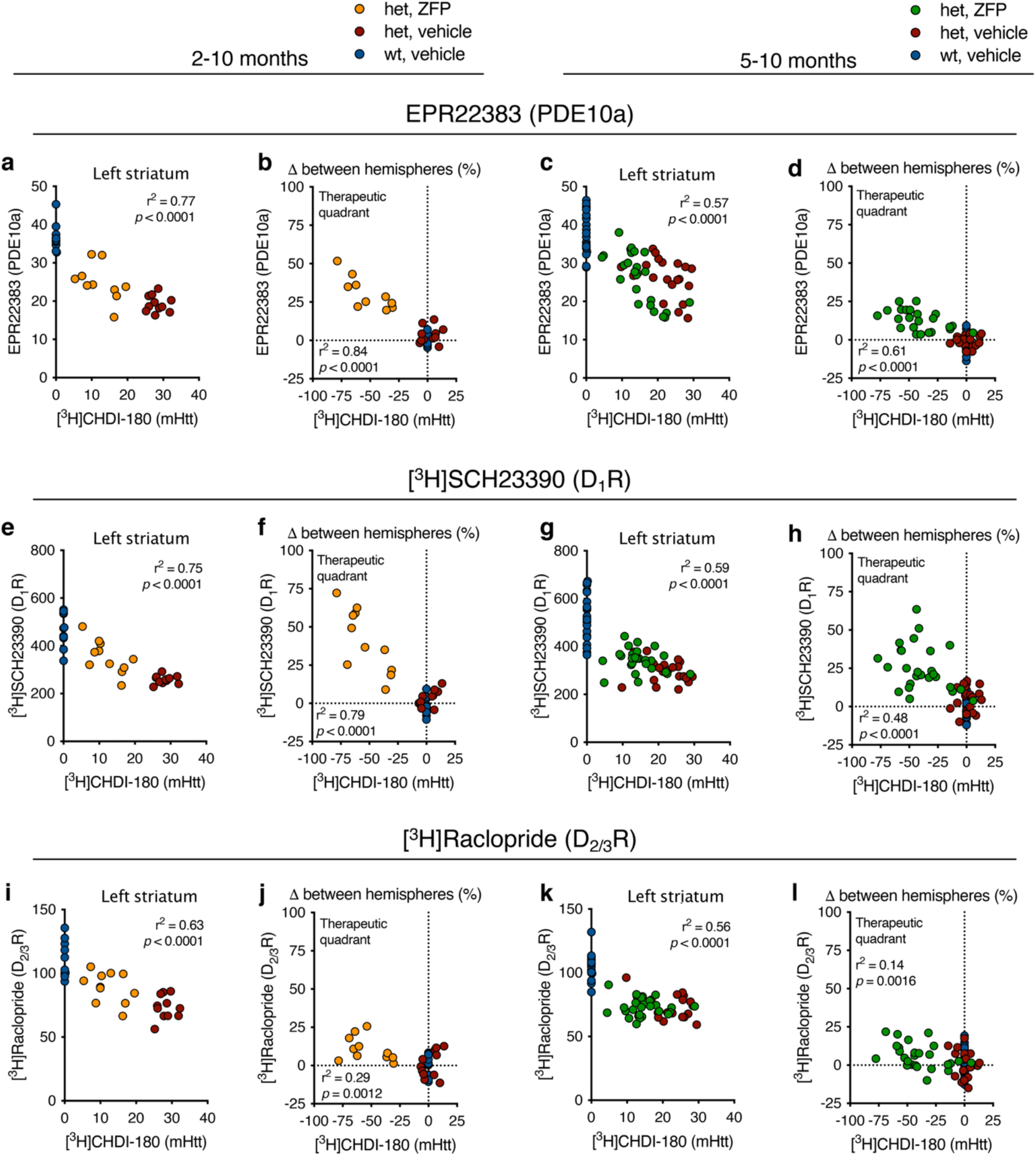
mHTT lowering following early and late ZFP intervention in zQ175DN mice is associated with preservation of striatal markers. *Post-mortem* correlation analyses between striatal specific binding of [³H]CHDI-180 (mHTT) in ZFP-treated hemisphere as well as percentage contralateral difference compared to PDE10a levels (**a**-**d**), D1R (**e**-**h**), and D2/3R (**i**-**l**) at 10 months of age following early (2 months) and late (5 months) intervention. Early intervention = het ZFP, n = 10; het vehicle, n = 11; wt vehicle, n = 11. Late intervention = het ZFP, n = 26; het vehicle, n = 20; wt vehicle, n = 23. Two-tailed Pearson correlation analyses, all points shown. Related to Figure 3 and 4.

**Extended Data Fig. 6.**
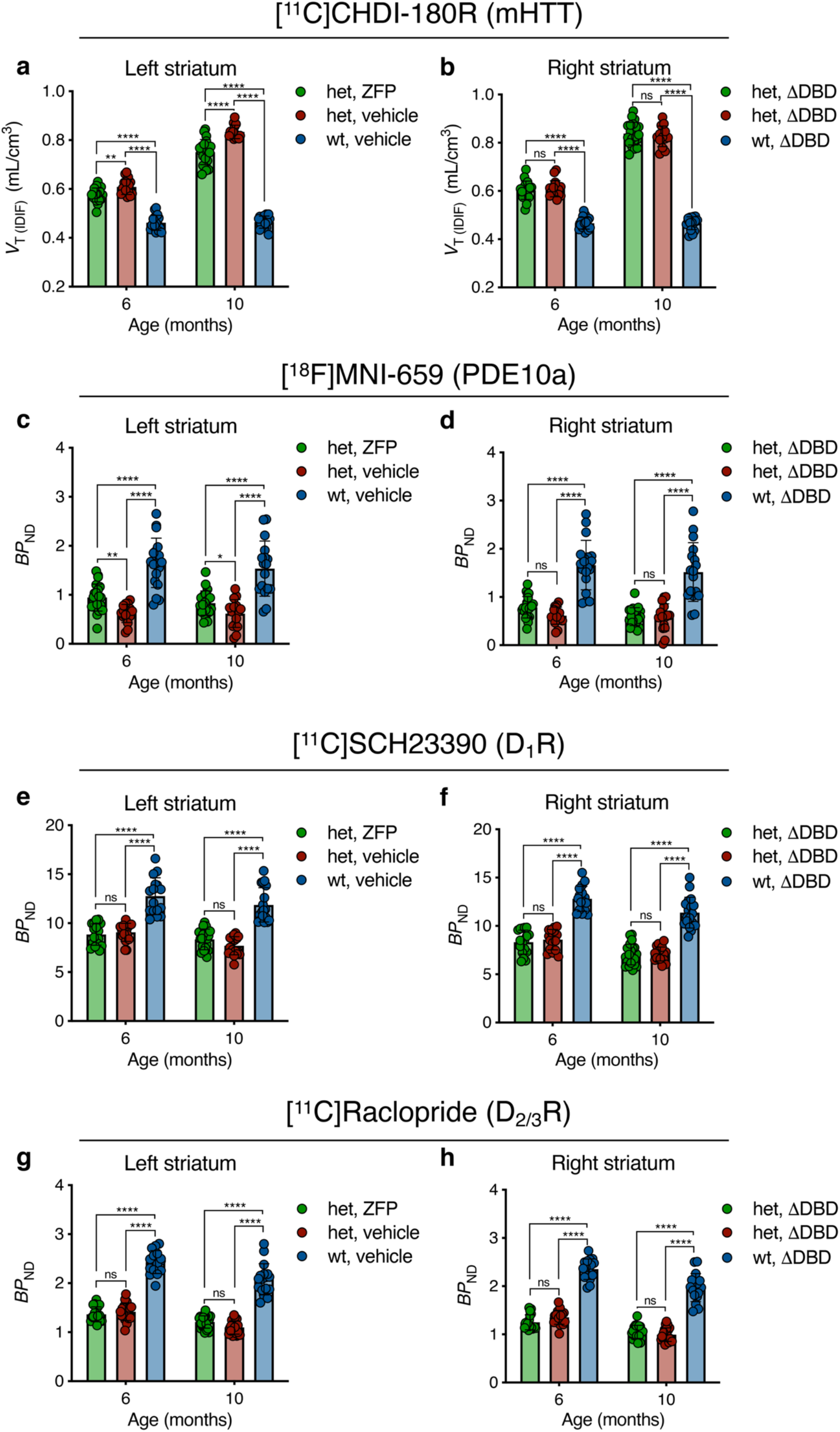
Longitudinal PET imaging quantification following ZFP intervention after mEM48-positive mHTT inclusions formation in the striatum of zQ175DN mice. **a**,**b**, Quantification of [^11^C]CHDI-180R *V*_T (IDIF)_ (mHTT inclusions) in left (**a**) and right (**b**) striatal hemispheres of zQ175DN wt vehicle, het vehicle, and het ZFP-treated mice at 6 months (het ZFP, n = 20; het vehicle, n = 17; wt vehicle, n = 17) and 10 months (het ZFP, n = 21; het vehicle, n = 19; wt vehicle, n = 18) of age following striatal injection at 5 months of age. Repeated measures with linear mixed model analysis with Tukey-Kramer correction; ** *P* < 0.01, **** *P* < 0.0001. Data are shown as mean ± s.d., all points shown. **c**,**d**, Quantification of [^18^F]MNI-659 *BP*_ND_ (PDE10a) in left (**c**) and right (**d**) striatal hemispheres of zQ175DN wt vehicle, het vehicle, and het ZFP-treated mice at 6 months (het ZFP, n = 20; het vehicle, n = 17; wt vehicle, n = 19) and 10 months (het ZFP, n = 21; het vehicle, n = 17; wt vehicle, n = 18) of age following striatal injection at 5 months of age. Repeated measures with linear mixed model analysis with Tukey-Kramer correction; * *P* < 0.05, ** *P* < 0.01, **** *P* < 0.0001. Data are shown as mean ± s.d., all points shown. **e**,**f**, Quantification of [^11^C]SCH23390 *BP*_ND_ (D_1_R) in left (**e**) and right (**f**) striatal hemispheres of zQ175DN wt vehicle, het vehicle, and het ZFP-treated mice at 6 months (het ZFP, n = 17; het vehicle, n = 16; wt vehicle, n = 16) and 10 months (het ZFP, n = 23; het vehicle, n = 20; wt vehicle, n = 19) of age following striatal injection at 5 months of age. Repeated measures with linear mixed model analysis with Tukey-Kramer correction; **** *P* < 0.0001. Data are shown as mean ± s.d., all points shown. **g**,**h**, Quantification of [^11^C]Raclopride *BP*_ND_ (D_2/3_R) in left (**g**) and right (**h**) striatal hemispheres of zQ175DN wt vehicle, het vehicle, and het ZFP-treated mice at 6 months (het ZFP, n = 17; het vehicle, n = 17; wt vehicle, n = 16) and 10 months (het ZFP, n = 23; het vehicle, n = 22; wt vehicle, n = 18) of age following striatal injection at 5 months of age. Repeated measures with linear mixed model analysis with Tukey-Kramer correction; **** *P* < 0.0001. Data are shown as mean ± s.d., all points shown. Related to Figure 4.

**Extended Data Fig. 7.**
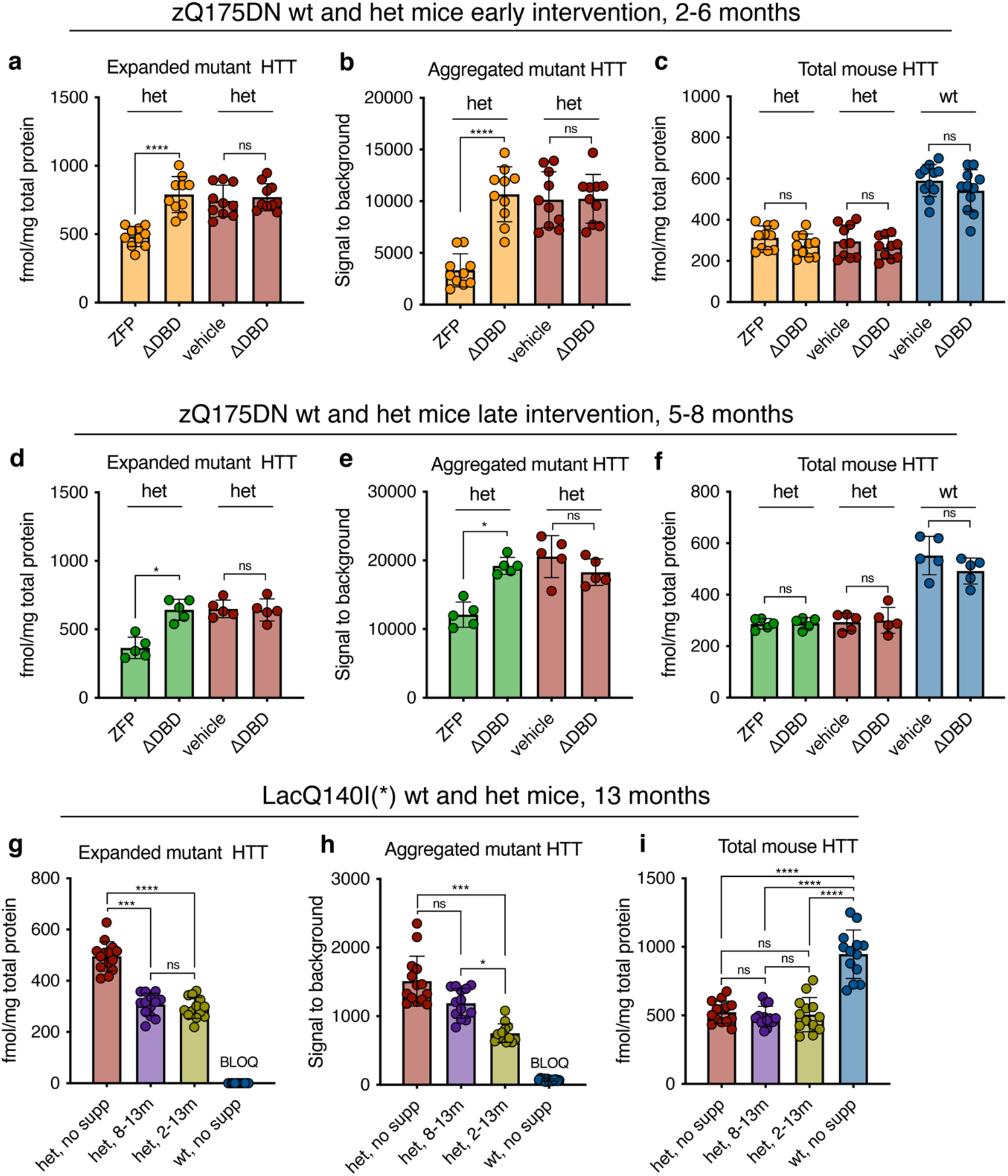
Quantification of wt Htt and mHTT of zQ175DN and LacQ140^I^(*) mice in mHTT lowering studies. **a-c**, Striatal quantification of expanded mHTT (**a**), aggregated mHTT (**b**), and wt Htt (**c**) in zQ175DN wt vehicle, het vehicle, and het ZFP-treated mice at 6 months (het ZFP, n = 11; het vehicle, n = 10; wt vehicle, n = 10) of age following striatal injection at 2 months of age. One-way ANOVA with Sidak’s multiple comparison test; **** *P* < 0.0001; Data are shown as mean ± s.d., all points are shown. **d-f**, Striatal quantification of expanded mHTT (**d**), aggregated mHTT (**e**), and wt Htt (**f**) in zQ175DN wt vehicle, het vehicle, and het ZFP-treated mice at 8 months (het ZFP, n = 11; het vehicle, n = 10; wt vehicle, n = 10) of age following striatal injection at 5 months of age. Kruskal-Wallis test with Dunn’s correction for multiple comparisons; * *P* < 0.05; Data are shown as mean ± s.d., all points are shown. **g-i**, Quantification of expanded mHTT (**g**), aggregated mHTT (**h**), and wt Htt (**i**) in the cerebellum of LacQ140^I^(*) mice at 13 months (het, no supp n = 14; het, 8-13m, n = 13; het, 2-13m, n = 13; wt, no supp, n = 13) of age. One-way ANOVA with Bonferroni’s multiple comparison test (g,i) and Kruskal-Wallis test with Dunn’s correction for multiple comparisons (h); * *P* < 0.05, *** *P* < 0.001, **** *P* < 0.0001; Data are shown as mean ± s.d., all points are shown. BLOQ, below limit of quantification.

**Extended Data Fig. 8.**
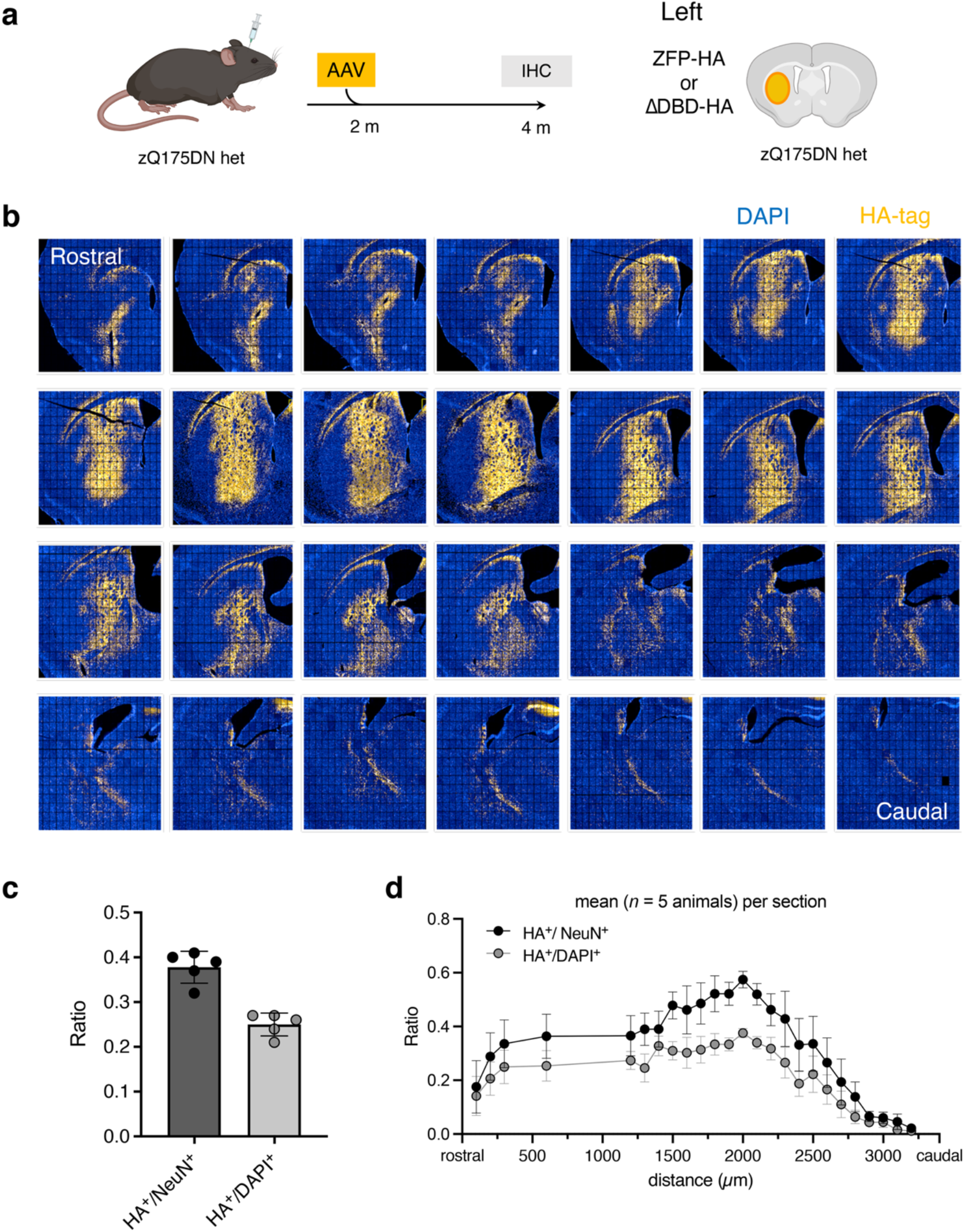
Biodistribution data for AAV ZFP in the striatum of zQ175DN mice. **a**, Experimental timeline for characterization of AAV ZFP and AAV ΔDBD biodistribution in zQ175DN mice. **b**,**c**, Representative striatal immunostaining for the neuronal transduction rate in the entire striatum (**b**) and resulting transduction rate following AAV ZFP and AAV ΔDBD biodistribution (**c**) as quantified with anti-HA. The mean transduction rate was 38%. Data are shown as mean ± s.d., all points are shown. **d**, Expression of the transgene is maximal close to the injection site and in dorsomedial portions of the striatum. Mean of all animals (*n* = 5) over all sections (*n* = 28-35). Data are shown as mean ± s.e.m. HA-tag, yellow; DAPI, blue.

**Extended Data Fig. 9.**
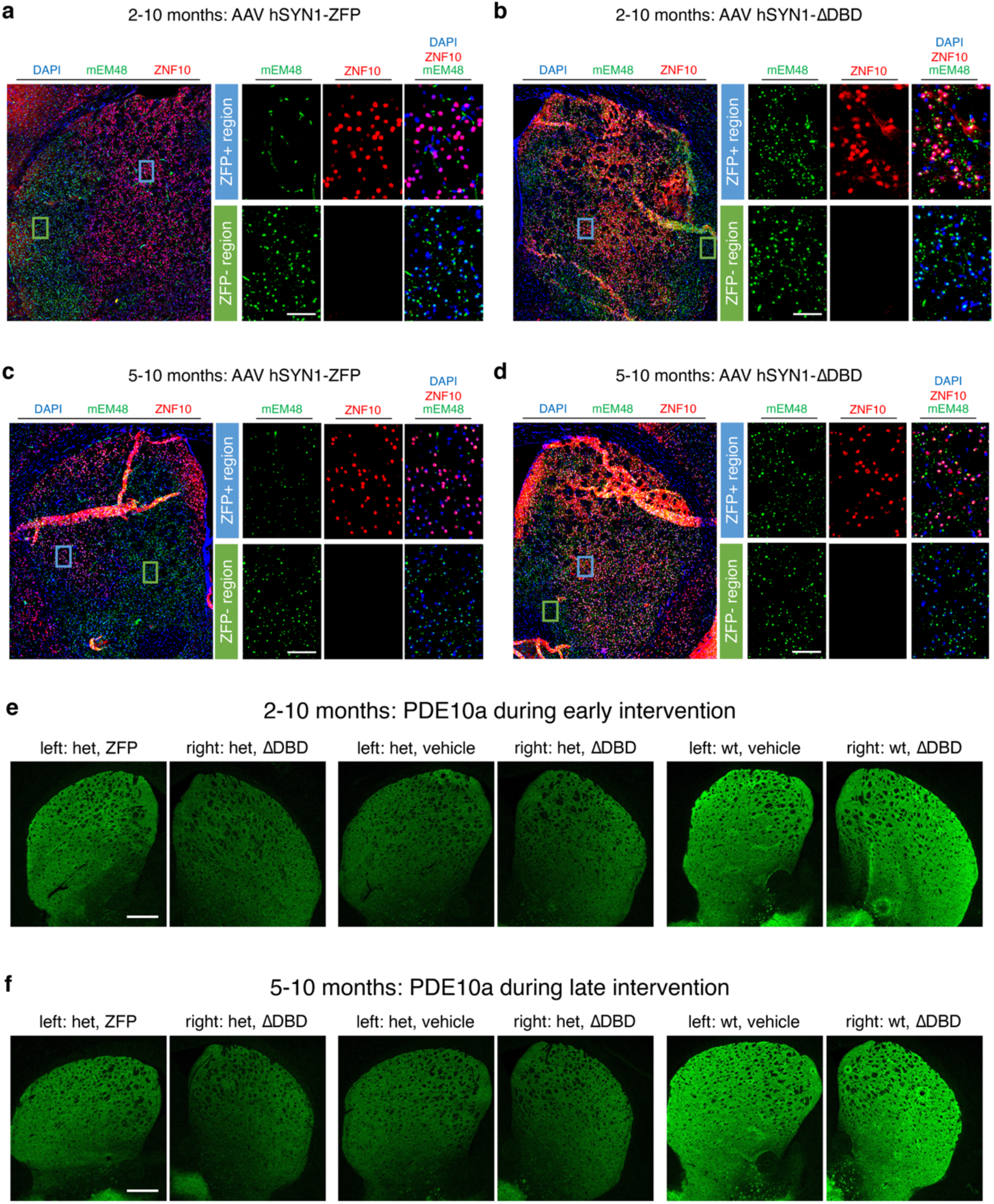
ZFP, mHTT aggregate, and PDE10a immunohistochemistry data for AAV ZFP treatment studies before or after mHTT inclusions formation in the striatum of zQ175DN mice. **a-d**, Representative striatal immunostaining images from 2-10 months zQ175DN cohorts (**a,b**) and 5-10 months zQ175DN cohorts (**c,d**) injected with AAV-delivered ZFP (left hemisphere) and ΔDBD (right hemisphere). Expanded ZFP^+^ and ZFP^−^ (**a,c**) as well as ΔDBD^+^ and ΔDBD^−^ (**b,d**) regions are shown at right. mHTT inclusions (mEM48), green; ZFP or ΔDBD (ZNF10), red; DAPI, blue. Scale bar, 50 *µ*m. **e,f** Representative striatal PDE10a immunostaining images from 2-10 months zQ175DN cohorts (**e**) and 5-10 months zQ175DN cohorts (**f**) injected with AAV-delivered ZFP (left hemisphere) and ΔDBD (right hemisphere). Scale bar, 500 *µ*m.

**Extended Data Fig. 10.**
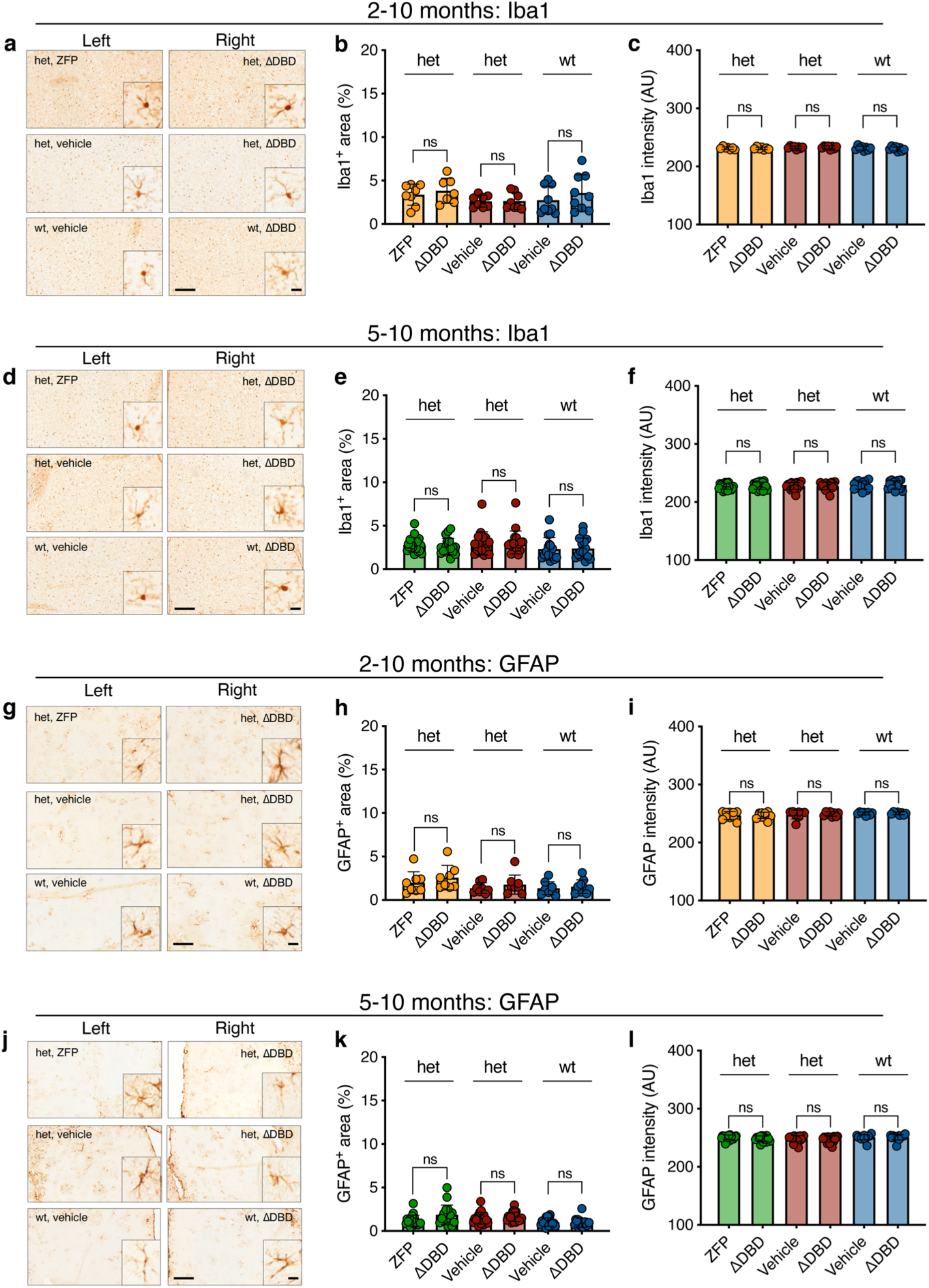
AAV-delivered ZFP before or after mEM48-positive mHTT inclusions formation in the striatum of zQ175DN mice does not induce gliosis. **a,d**, Representative striatal Iba1 immunostaining images from 2-10 months zQ175DN cohorts (**a**) and 5-10 months zQ175DN cohorts (**d**) injected with AAV-delivered ZFP, ΔDBD, or vehicle. Scale bar, 200 *µ*m, inset 20 *µ*m. **b,c**, Quantification of Iba1+ area (**b**) and Iba1 intensity (**c**) in the striatum of zQ175DN wt vehicle, het vehicle, and het ZFP-treated mice at 10 months (het ZFP, n = 7; het vehicle, n = 8; wt vehicle, n = 9) of age following striatal injection at 2 months of age. Two-way ANOVA with Bonferroni’s multiple comparison test; mean ± s.d., all points are shown. **e,f**, Quantification of Iba1+ area (**e**) and Iba1 intensity (**f**) in the striatum of zQ175DN wt vehicle, het vehicle, and het ZFP-treated mice at 10 months (het ZFP, n = 19; het vehicle, n = 16; wt vehicle, n = 16) of age following striatal injection at 5 months of age. Two-way ANOVA with Bonferroni’s multiple comparison test; mean ± s.d., all points are shown. **g,j**, Representative striatal GFAP immunostaining images from 2-10 months zQ175DN cohorts (**g**) and 5-10 months zQ175DN cohorts (**j**) injected with AAV-delivered ZFP, ΔDBD, or vehicle. S Scale bar, 200 *µ*m, inset 20 *µ*m. **h,i**, Quantification of GFAP+ area (**h**) and GFAP intensity (**i**) in the striatum of zQ175DN wt vehicle, het vehicle, and het ZFP-treated mice at 10 months (het ZFP, n = 8; het vehicle, n = 8; wt vehicle, n = 8) of age following striatal injection at 2 months of age. Two-way ANOVA with Bonferroni’s multiple comparison test; mean ± s.d., all points are shown. **k,l**, Quantification of GFAP+ area (**k**) and GFAP intensity (**l**) in the striatum of zQ175DN wt vehicle, het vehicle, and het ZFP-treated mice at 10 months (het ZFP, n = 18; het vehicle, n = 19; wt vehicle, n = 16) of age following striatal injection at 5 months of age. Two-way ANOVA with Bonferroni’s multiple comparison test; mean ± s.d., all points are shown.

**Supplementary Table 1.**
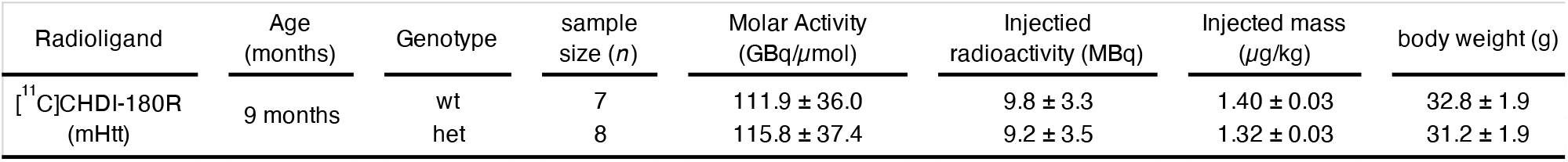
Sample size, injected radioactivity, injected mass, and bodyweight of zQ175DN mice imaged during the [^11^C]CHDI-180R radioligand validation study. Values are expressed as mean ± SD.

| Radioligand | Age (months) | Genotype | sample size (n) | Molar Activity (GBq/μmol) | Injected radioactivity (MBq) | Injected mass (μg/kg) | body weight (g) |
| --- | --- | --- | --- | --- | --- | --- | --- |
| <sup>11</sup>C]CHDI-180R (mHtt) | 9 months | wt | 7 | 111.9 ± 36.0 | 9.8 ± 3.3 | 1.40 ± 0.03 | 32.8 ± 1.9 |
|  |  | het | 8 | 115.8 ± 37.4 | 9.2 ± 3.5 | 1.32 ± 0.03 | 31.2 ± 1.9 |

**Supplementary Table 2.** Kinetic parameters for [^11^C]CHDI-180R determined using 2-tissue compartment model and Logan graphical analysis in het zQ175DN. Values are expressed as mean ± SD.

| Brain region | $K_1$<br>(mL/cm<sup>3</sup>/min) | $k_2$<br>(per min) | $k_3$<br>(per min) | $k_4$<br>(per min) | $V_T$ (IDIF) (2TCM)<br>(mL/cm<sup>3</sup>) | $V_T$ (IDIF) (Logan)<br>(mL/cm<sup>3</sup>) |
| --- | --- | --- | --- | --- | --- | --- |
| Striatum | 0.97 ± 0.07 | 1.83 ± 0.14 | 0.048 ± 0.012 | 0.085 ± 0.018 | 0.83 ± 0.05 | 0.82 ± 0.04 |
| Motor cortex | 1.01 ± 0.07 | 2.17 ± 0.25 | 0.063 ± 0.020 | 0.088 ± 0.021 | 0.80 ± 0.05 | 0.79 ± 0.04 |
| Hippocampus | 1.12 ± 0.15 | 2.19 ± 0.24 | 0.048 ± 0.011 | 0.097 ± 0.020 | 0.79 ± 0.05 | 0.78 ± 0.04 |
| Cerebellum | 1.18 ± 0.10 | 2.70 ± 0.32 | 0.090 ± 0.041 | 0.159 ± 0.054 | 0.68 ± 0.05 | 0.69 ± 0.04 |

**Supplementary Table 3.** Sample size, injected radioactivity, injected mass, and bodyweight of zQ175DN mice imaged the [^11^C]CHDI-180R longitudinal study. Values are expressed as mean ± SD.

| Radioligand | Age (months) | Genotype | sample size (n) | Molar Activity (GBq/μmol) | Injected radioactivity (MBq) | Injected mass (μg/kg) | body weight (g) |
| --- | --- | --- | --- | --- | --- | --- | --- |
| <sup>11</sup>C]CHDI-180R (mHtt) | 3 months | wt | 19 | 140.1 ± 33.8 | 5.6 ± 1.6 | 1.11 ± 0.07 | 27.4 ± 1.6 |
|  |  | het | 21 | 153.9 ± 38.6 | 5.9 ± 1.6 | 1.12 ± 0.07 | 26.6 ± 1.3 |
| <sup>11</sup>C]CHDI-180R (mHtt) | 6 months | wt | 15 | 172.3 ± 64.2 | 7.8 ± 3.0 | 0.98 ± 0.10 | 32.0 ± 2.8 |
|  |  | het | 23 | 180.7 ± 59.2 | 7.7 ± 3.1 | 0.99 ± 0.08 | 29.3 ± 1.5 |
| <sup>11</sup>C]CHDI-180R (mHtt) | 9 months | wt | 13 | 192.1 ± 90.7 | 8.1 ± 3.9 | 0.99 ± 0.06 | 32.8 ± 2.7 |
|  |  | het | 20 | 192.0 ± 73.3 | 7.0 ± 2.5 | 1.02 ± 0.06 | 27.6 ± 1.8 |
| <sup>11</sup>C]CHDI-180R (mHtt) | 13 months | wt | 12 | 94.3 ± 18.7 | 5.4 ± 0.9 | 1.08 ± 0.09 | 34.8 ± 2.8 |
|  |  | het | 17 | 102.8 ± 15.6 | 4.1 ± 0.5 | 1.03 ± 0.16 | 26.3 ± 1.5 |

**Supplementary Table 4.** Sample size calculations at desired therapeutic effects for the design of disease-modifying interventions using striatal [11C]CHDI-180R PET imaging as endpoint. Values are determined based on a one-tailed test, with *α* = 0.05 and power (1-*β*) = 0.80.

| Therapeutic effect (%) | Sample size required per experimental arm (*n*) |  |  |  |
| --- | --- | --- | --- | --- |
|  | 3 months | 6 months | 9 months | 13 months |
| 100% | *n* = 12 | *n* = 3 | *n* = 3 | *n* = 3 |
| 90% | *n* = 15 | *n* = 3 | *n* = 3 | *n* = 3 |
| 80% | *n* = 18 | *n* = 3 | *n* = 3 | *n* = 3 |
| 70% | *n* = 23 | *n* = 3 | *n* = 3 | *n* = 3 |
| 60% | *n* = 31 | *n* = 3 | *n* = 3 | *n* = 3 |
| 50% | *n* = 44 | *n* = 4 | *n* = 3 | *n* = 3 |
| 40% | *n* = 66 | *n* = 6 | *n* = 3 | *n* = 3 |
| 30% | *n* = 113 | *n* = 8 | *n* = 4 | *n* = 3 |
| 20% | *n* = 258 | *n* = 16 | *n* = 6 | *n* = 4 |

**Supplementary Table 5.** Sample size, injected radioactivity, injected mass, and bodyweight of zQ175DN mice imaged during the longitudinal mHtt-ZFP2M treatment study. Values are expressed as mean ± SD.

| Radioligand | Age (months) | Genotype | Condition | sample size (n) | Molar Activity (GBq/μmol) | Injected radioactivity (MBq) | Injected mass (μg/kg) | body weight (g) |
| --- | --- | --- | --- | --- | --- | --- | --- | --- |
| $[^{11}\text{C}]\text{CHDI-180R}$ (mHtt) | 3 months | het | ZFP | 21 | 128.6 ± 20.2 | 3.4 ± 0.5 | 0.73 ± 0.13 | 24.8 ± 1.4 |
| | | het | vehicle | 21 | 131.2 ± 21.3 | 3.5 ± 0.7 | 0.73 ± 0.11 | 25.3 ± 0.9 |
| | | wt | vehicle | 20 | 135.5 ± 26.6 | 3.6 ± 0.6 | 0.71 ± 0.10 | 25.3 ± 1.7 |
| $[^{18}\text{F}]\text{MNI-659}$ (PDE10a) | 3 months | het | ZFP | 21 | 260.6 ± 6.1 | 5.4 ± 1.9 | 0.65 ± 0.20 | 24.7 ± 0.2 |
| | | het | vehicle | 22 | 264.7 ± 60.3 | 5.1 ± 1.9 | 0.52 ± 0.22 | 25.3 ± 1.0 |
| | | wt | vehicle | 20 | 271.2 ± 52.6 | 4.9 ± 1.8 | 0.61 ± 0.17 | 25.6 ± 1.5 |
| $[^{11}\text{C}]\text{CHDI-180R}$ (mHtt) | 6 months | het | ZFP | 18 | 127.7 ± 21.3 | 4.3 ± 0.8 | 0.85 ± 0.11 | 28.7 ± 1.4 |
| | | het | vehicle | 19 | 125.7 ± 19.4 | 4.2 ± 0.8 | 0.86 ± 0.11 | 28.6 ± 1.4 |
| | | wt | vehicle | 19 | 125.2 ± 20.5 | 4.3 ± 0.7 | 0.83 ± 0.10 | 30.3 ± 2.5 |
| $[^{18}\text{F}]\text{MNI-659}$ (PDE10a) | 6 months | het | ZFP | 21 | 697.9 ± 158.0 | 8.7 ± 1.9 | 0.42 ± 0.18 | 28.2 ± 1.5 |
| | | het | vehicle | 21 | 664.1 ± 119.2 | 8.8 ± 1.4 | 0.42 ± 0.16 | 28.7 ± 1.3 |
| | | wt | vehicle | 18 | 708.3 ± 129.1 | 8.9 ± 1.9 | 0.43 ± 0.19 | 30.0 ± 2.6 |
| $[^{11}\text{C}]\text{CHDI-180R}$ (mHtt) | 10 months | het | ZFP | 18 | 161.1 ± 27.6 | 4.8 ± 0.8 | 0.78 ± 0.08 | 28.0 ± 1.8 |
| | | het | vehicle | 19 | 152.6 ± 20.8 | 4.8 ± 0.8 | 0.81 ± 0.08 | 28.2 ± 1.7 |
| | | wt | vehicle | 19 | 151.3 ± 27.8 | 5.5 ± 1.3 | 0.79 ± 0.06 | 33.5 ± 3.8 |
| $[^{18}\text{F}]\text{MNI-659}$ (PDE10a) | 10 months | het | ZFP | 20 | 716.2 ± 232.3 | 8.9 ± 1.8 | 0.41 ± 0.16 | 28.1 ± 2.0 |
| | | het | vehicle | 20 | 732.6 ± 199.4 | 8.6 ± 1.6 | 0.38 ± 0.16 | 28.1 ± 1.6 |
| | | wt | vehicle | 18 | 780.7 ± 222.3 | 9.1 ± 1.6 | 0.40 ± 0.22 | 33.0 ± 3.8 |

**Supplementary Table 6.**
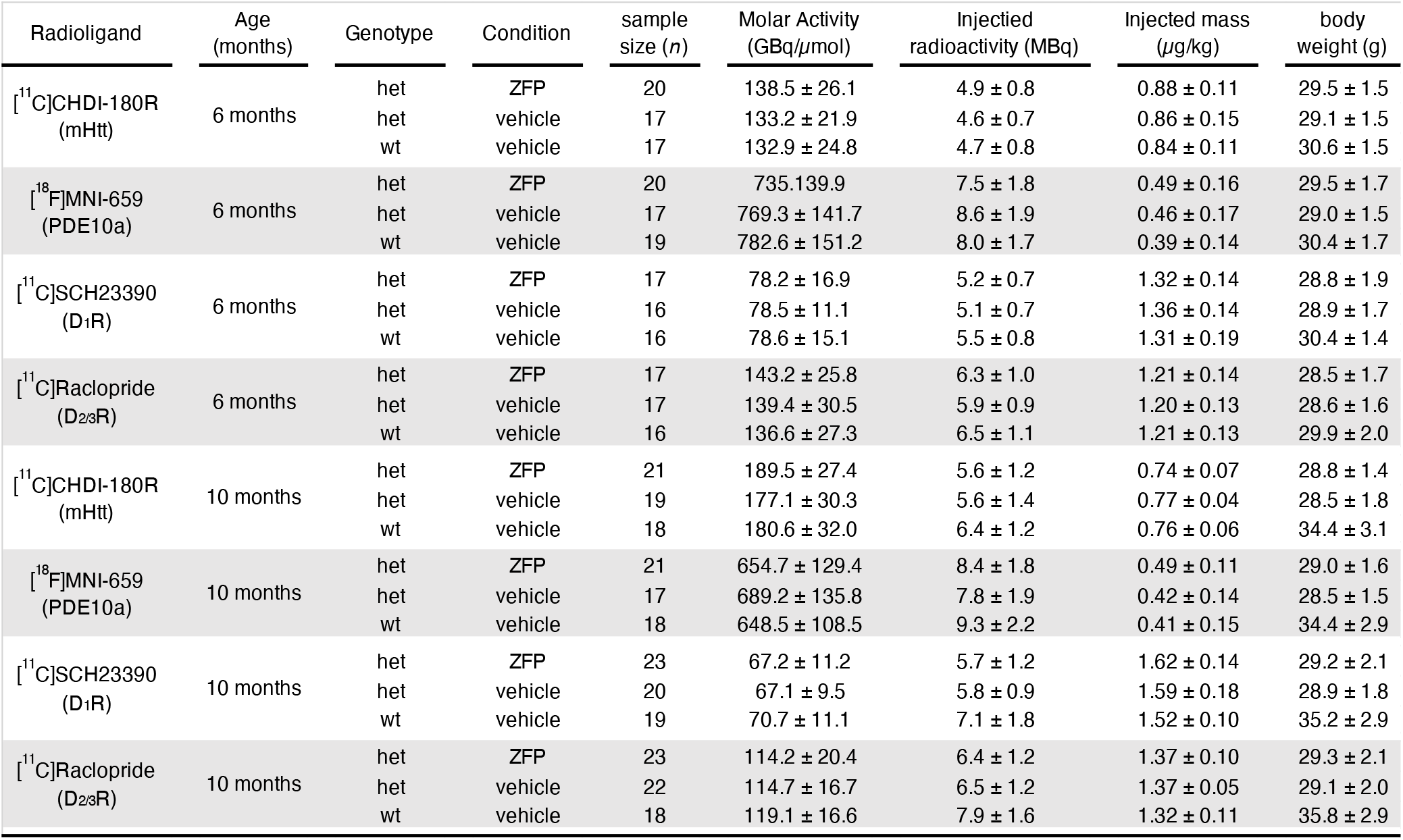
Sample size, injected radioactivity, injected mass, and bodyweight of zQ175DN mice imaged during the longitudinal mHtt-ZFP5M treatment study. Values are expressed as mean ± SD.

**Supplementary Table 7.** Sample size, injected radioactivity, injected mass and bodyweight of LacQ140I(*) mice imaged with [^11^C]CHDI-180R. Values are expressed as mean ± SD.

| Radioligand | Age (months) | Genotype | Condition | sample size (n) | Molar Activity (GBq/μmol) | Injected radioactivity (MBq) | Injected mass (μg/kg) | body weight (g) |
| --- | --- | --- | --- | --- | --- | --- | --- | --- |
| <sup>11</sup>C]CHDI-180R (mHtt) | 13 months | wt | no suppression | 13 | 176.4 ± 40.9 | 6.3 ± 1.4 | 0.77 ± 0.11 | 35.4 ± 2.6 |
|  |  | het | no suppression | 13 | 184.7 ± 44.8 | 5.9 ± 1.2 | 0.75 ± 0.11 | 32.7 ± 2.7 |
|  |  | het | 2-13m suppression | 12 | 161.4 ± 45.5 | 5.2 ± 1.1 | 0.78 ± 0.14 | 33.2 ± 1.4 |
|  |  | het | 8-13m suppression | 14 | 176.8 ± 46.2 | 6.0 ± 1.3 | 0.79 ± 0.20 | 32.2 ± 2.3 |

**Supplementary Table 8.** Contralateral quantification in zQ175DN mice imaged during the longitudinal mHtt-ZFP2M treatment study. Difference between left and right striatum calculated using a within-subject two-paired t-test. Values are expressed as mean ± SD.

| Radioligand | Time period (months) | Genotype | Treatment (left vs. right striatum) | sample size (n) | Endpoint | Left striatum (mean ± SD) | Right striatum (mean ± SD) | % Difference Left vs. Right striatum | Within subject paired t-test |
| --- | --- | --- | --- | --- | --- | --- | --- | --- | --- |
| $[^{11}\text{C}]\text{CHDI-180R}$ (mHtt) | 2-3 months | het | ZFP vs $\Delta\text{DBD}$ | 21 | $V_{\text{T}}$ (IDIF) | 0.50 ± 0.03 | 0.51 ± 0.03 | -2.8% | * $P = 0.0122$ |
| | | het | vehicle vs $\Delta\text{DBD}$ | 21 | | 0.50 ± 0.03 | 0.50 ± 0.03 | 0.0% | ns |
| | | wt | vehicle vs $\Delta\text{DBD}$ | 20 | | 0.47 ± 0.02 | 0.47 ± 0.02 | -0.1% | ns |
| $[^{18}\text{F}]\text{MNI-659}$ (PDE10a) | 2-3 months | het | ZFP vs $\Delta\text{DBD}$ | 21 | $BP_{\text{ND}}$ | 1.50 ± 0.45 | 1.23 ± 0.35 | 22.7% | **** $P < 0.0001$ |
| | | het | vehicle vs $\Delta\text{DBD}$ | 22 | | 1.25 ± 0.37 | 1.22 ± 0.35 | 1.4% | ns |
| | | wt | vehicle vs $\Delta\text{DBD}$ | 20 | | 1.94 ± 0.49 | 1.90 ± 0.53 | 2.7% | ns |
| $[^{11}\text{C}]\text{CHDI-180R}$ (mHtt) | 2-6 months | het | ZFP vs $\Delta\text{DBD}$ | 18 | $V_{\text{T}}$ (IDIF) | 0.53 ± 0.03 | 0.58 ± 0.03 | -9.0% | **** $P < 0.0001$ |
| | | het | vehicle vs $\Delta\text{DBD}$ | 19 | | 0.59 ± 0.03 | 0.59 ± 0.03 | 0.1% | ns |
| | | wt | vehicle vs $\Delta\text{DBD}$ | 19 | | 0.45 ± 0.02 | 0.45 ± 0.02 | -1.2% | ns |
| $[^{18}\text{F}]\text{MNI-659}$ (PDE10a) | 2-6 months | het | ZFP vs $\Delta\text{DBD}$ | 21 | $BP_{\text{ND}}$ | 0.99 ± 0.45 | 0.54 ± 0.25 | 98.1% | **** $P < 0.0001$ |
| | | het | vehicle vs $\Delta\text{DBD}$ | 21 | | 0.59 ± 0.19 | 0.56 ± 0.18 | 4.3% | ns |
| | | wt | vehicle vs $\Delta\text{DBD}$ | 18 | | 1.65 ± 0.43 | 1.67 ± 0.48 | -1.2% | ns |
| $[^{11}\text{C}]\text{CHDI-180R}$ (mHtt) | 2-10 months | het | ZFP vs $\Delta\text{DBD}$ | 18 | $V_{\text{T}}$ (IDIF) | 0.66 ± 0.04 | 0.79 ± 0.06 | -16.3% | **** $P < 0.0001$ |
| | | het | vehicle vs $\Delta\text{DBD}$ | 19 | | 0.80 ± 0.05 | 0.79 ± 0.05 | 2.1% | ns |
| | | wt | vehicle vs $\Delta\text{DBD}$ | 19 | | 0.45 ± 0.02 | 0.45 ± 0.02 | -0.8% | ns |
| $[^{18}\text{F}]\text{MNI-659}$ (PDE10a) | 2-10 months | het | ZFP vs $\Delta\text{DBD}$ | 20 | $BP_{\text{ND}}$ | 0.99 ± 0.49 | 0.54 ± 0.30 | 98.1% | **** $P < 0.0001$ |
| | | het | vehicle vs $\Delta\text{DBD}$ | 20 | | 0.54 ± 0.27 | 0.54 ± 0.29 | 2.0% | ns |
| | | wt | vehicle vs $\Delta\text{DBD}$ | 18 | | 1.56 ± 0.57 | 1.59 ± 0.60 | -1.7% | ns |

**Supplementary Table 9.**
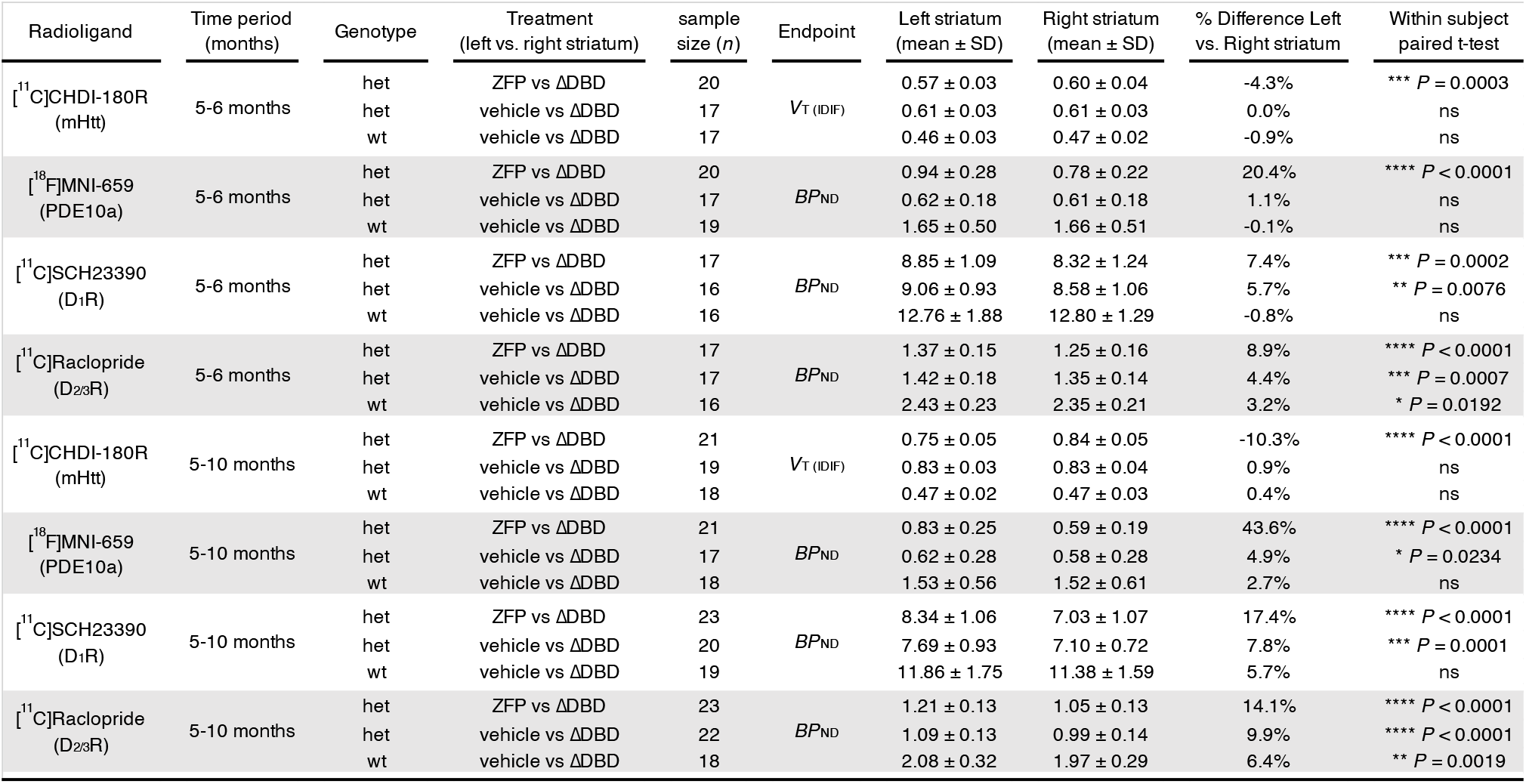
Contralateral quantification in zQ175DN mice imaged during the longitudinal mHtt-ZFP5M treatment study. Difference between left and right striatum calculated using a within-subject two-paired t-test. Values are expressed as mean ± SD.

**Supplementary Table 10.**
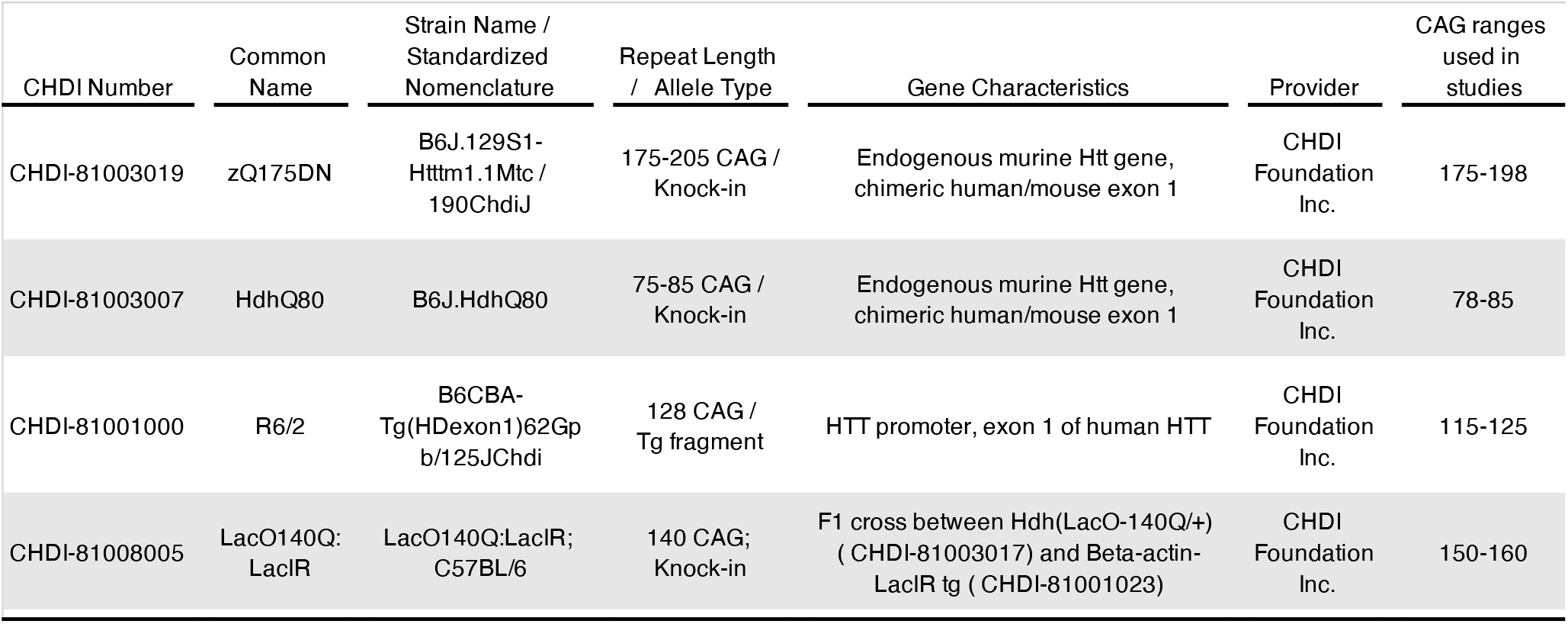
Mouse models used for studies.

**Supplementary Table 11.** Human brain tissue used for autoradiography and histological analysis. Demographic information for human HD and control brains. Abbreviations: F = female, HD = Huntington’s disease, M = male, n/a = not available, NFT = neurofibrillary tangles, PMI = postmortem interval. Tissue was obtained from Netherland Brain Bank (NBB).

| ID | Diagnosis | Gender | Age at death (Years) | PMI (hours) | CAG repeats | Vonsattel grade | Braak stage (NFT) |
| --- | --- | --- | --- | --- | --- | --- | --- |
| 2014-043 | Control | F | 60 | 8 | n/a | n/a | 0 |
| 2017-005 | Control | F | 60 | 5.5 | n/a | n/a | 0 |
| 2017-060 | HD | F | 57 | 6.5 | n/a | n/a | 0 |
| 2008-044 | HD | M | 59 | 5 | n/a | n/a | n/a |

